# Major transitions in early coral development: novel insights enabled by visualisation of a comprehensive transcriptomic dataset for *Acropora millepora*

**DOI:** 10.1101/2025.06.16.660037

**Authors:** Ramona Brunner, Mila Grinblat, Aurelie Moya, Sylvain Foret, David C Hayward, Bruno Lapeyre, Eldon E Ball, Ira Cooke, David J Miller

**Affiliations:** ARC Centre of Excellence for Coral Reef Studies, James Cook University, Townsville, QLD 4811 Australia; Molecular Genetics Unit, James Cook University, Townsville, QLD 4811 Australia; Research School of Biology, Australian National University, Acton, ACT 2601 Australia; Laboratoire d’excellence CORAIL, Centre de Recherches Insulaires et Observatoire de l’Environnement (CRIOBE), Moorea, B.P. 1013, Papeete, French Polynesia; Department of Marine Biology and Aquaculture, James Cook University, Townsville, QLD 4811 Australia

**Keywords:** Maternal-zygotic transition, juvenile-adult transition, metamorphosis, coral development, larval settlement

## Abstract

**Background:** Given the ecological importance of reef-building corals (Scleractinia), it is perhaps surprising that the molecular mechanisms underlying many of the morphological and metabolic changes during their development remain unclear. In part, this is due to the lack of a comprehensive transcriptomic dataset for any coral. A second challenge in the analysis of such non-model developmental datasets is that the volume of data often complicates its interpretation.

**Results:** To overcome these limitations, we profiled gene expression in *Acropora millepora* across 26 life stages from unfertilised eggs to juvenile polyps and developed an interactive online tool based on the R-application Shiny to simultaneously visualise changes in the expression of large numbers of genes. As expected, major transcriptomic changes (transitions) occurred during gastrulation and the acquisition of competence. Surprisingly, however, settlement triggered by using the natural inducer CCA did not immediately lead to major changes in gene expression, but a major transition involving many genes was observed 3 - 6 hours after settlement induction.

**Conclusions:** We hope that providing access to this extensive developmental transcriptome dataset and software to facilitate its analysis will expedite a better understanding of the changes that occur during coral development. The online tool is available at https://amil-deview.mmb.group.

## Background

Broadcast spawning corals, such as *Acropora*, release buoyant eggs and sperm and fertilisation occurs in the water column. The early developmental stages are planktonic, ultimately reaching a motile stage known as a planula larva which becomes “competent” – that is, it becomes capable of settlement and metamorphosis - after a period of development of variable duration (Figure 1).

**Figure 1.**
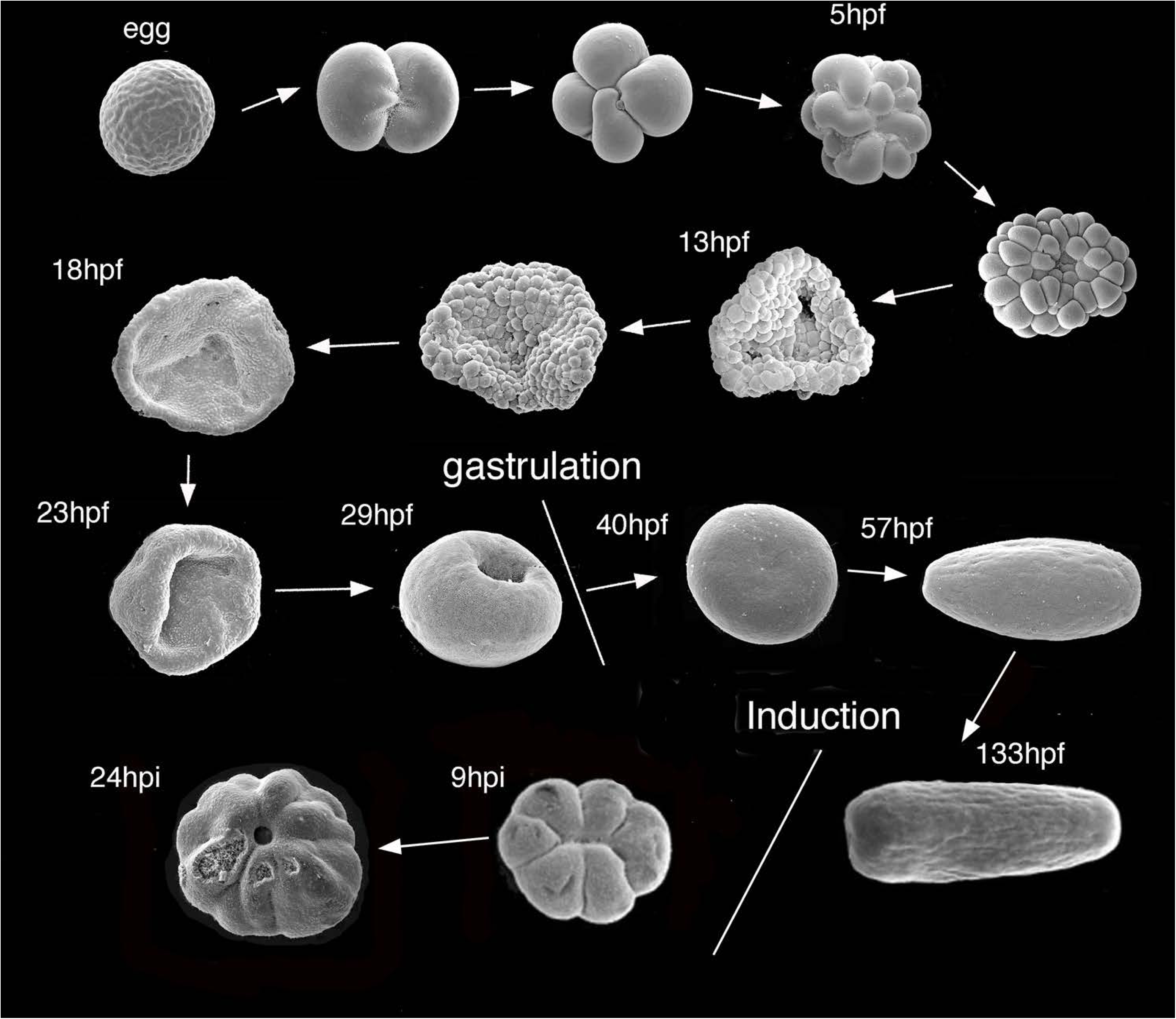
Images showing the morphologies recorded at the time of collection at the timepoints indicated. Times are given in hours post fertilisation (hpf) and hours post induction (hpi). The places in the developmental sequence of two major events, gastrulation and induction with a settlement cue, are indicated. Some of the SEM images have been previously published in Grasso et al. 2008, Hayward et al. 2011, and Ball et al. 2021.

Competence is marked by a change in behaviour, from horizontal swimming to swimming into the substratum and testing it for chemical cues. In nature, the settlement of competent larvae generally involves selection of appropriate sites, based on perception of a range of cues that include small molecules, at least some of which are produced by bacteria associated with substrate biofilms and algae (e.g. Tebben et al. 2015; Siboni et al. 2020; Quinlan et al. 2023; Turnlund et al. 2025). The quality and nature of available light also plays a part in settlement site selection in the case of many reef-building corals (Mason and Cohen, 2012; Strader et al. 2015; Mason et al. 2023).

Various aspects of early coral development have previously been studied (Table 1), with the initial focus being characterisation of individual genes homologous to those playing key developmental roles in bilaterians (Ball et al. 2002; Hayward et al. 2002; Ball et al. 2004; Hayward et al. 2004). The first larger scale analyses were based on microarrays (Grasso et al. 2008; Reyes-Bermudez et al. 2009; Grasso et al. 2011), 454 GSFlx sequencing (Meyer et al. 2009), SOLiD sequencing (Meyer et al. 2011) or subtractive hybridisation (Hayward et al. 2011). More recent studies have used high-throughput DNA sequencing technologies to characterise specific developmental stages (Summarised in Table 1). Much of this work has focussed on members of the Acroporidae (genera Acropora and Montipora), with sampling regimes designed to address specific questions. For example, Chille et al. (2021) concentrated on gene expression changes *in Montipora digitata* during the maternal/zygotic transition (MZT). However, two recent developmental studies on *M. digitata* have gone beyond transcriptomics and are particularly relevant to the work presented here. He et al. (2024) combined spatial gene expression data with a comprehensive temporal dataset to study the segmentation of the gastric cavity, and Huffmeyer et al. (2025) supplemented transcriptomic data with metabolomic analyses to investigate the nutritional shift from maternal lipids to symbiont photosynthates in this vertically transmitting coral. Much higher cellular resolution expression datasets are becoming available for scleractinians (Levy et al. 2021; Gilbert et al. 2024; Ramon-Mateu et al. 2025), but so far these have been limited to a few developmental stages.

**Table 1.**
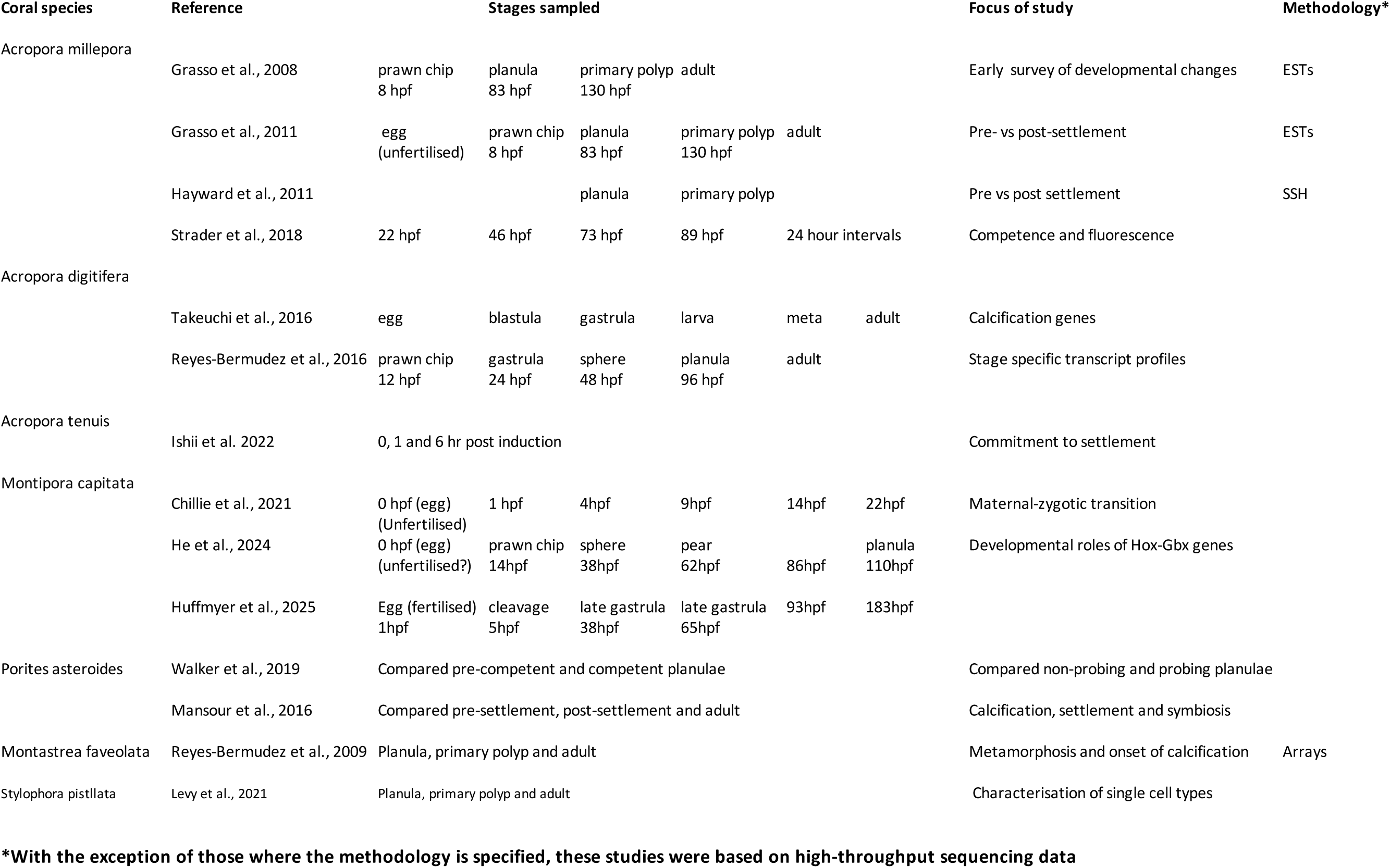
Previous studies on the molecular bases of early development in hard corals.

To address the need for a comprehensive transcriptomic dataset, we profiled gene expression of *Acropora millepora* across 26 life stages from fertilised eggs to juvenile polyps and developed an interactive online tool based on the Shiny web framework to visualise changes in the expression of groups of genes. We present an overview of the data, focussing on major transitions in gene expression and on processes not previously investigated, but hope that further exploration of the dataset via the website facilitates progress towards solutions for the crises currently faced by coral reefs worldwide. Our study also reveals the importance of combining spatial expression data, as revealed by in situ hybridisation, with the data obtained by RNAseq, in order to better understand gene function.

## Materials and Methods

Colonies of *Acropora millepora* were collected at Orpheus Island, Queensland, Australia, (18.6141° S, 146.4892° E) under GBRMPA permit G13/36196.1 shortly before they spawned at approximately 9pm on 21 November 2013. The samples sequenced were the result of crosses between two colonies and were raised in three separate flow-through tanks to control for tank effects.

Material was collected from 26 different time points of which 14 were sampled before and 12 after settlement induction. Pre-settlement, samples were collected from unfertilised eggs, at 5 hours post-fertilisation (hpf; 16-64 cell stages), 13hpf (prawnchip), 18hpf, 23hpf, 29hpf, 35hpf, 40hpf (sphere, some swimming), 45hpf (spheres, all swimming), 57hpf, 69hpf, 81hpf (planulae prior to the searching phase), 93hpf (searching planulae) and at 133hpf. At these stages, approx. 500 embryos per replicate (n=3) and cross (n = 2) were immersed in 2 mL ethanol (100%). This caused them to sink to the tube bottom, so that the overlying liquid was removed before tubes were snap-frozen in liquid nitrogen.

To assess settlement competency, between 69 and 93hpf, aliquots of planulae were exposed (independently) to either ethanolic extracts of the crustose coralline alga (CCA) *Porolithon onkodes*, (kindly provided by Mikhail V. Matz) or the neuropeptide Hym-248, both of which are known to induce settlement of *Acropora* planulae. At the 93 hpf stage 100% of planulae rapidly responded to the CCA extract by undergoing settlement followed by metamorphosis, hence the settlement-induced stages used for RNAseq were derived from planulae exposed to this inducer. Post-settlement samples were collected by stocking 144 petri dishes (2 replicates x 3 larval tanks x 2 crosses x 12 time points) with respectively 100 larvae at the age of 133 h. Settlement was induced with ethanolic CCA extract in all petri dishes at the same time. At the respective sampling times of 15 minutes after induction (mpi), 30mpi, 1, 2, 3, 6, 9, 12, 18, 24, 48 and 72 hours post induction (hpi), unattached larvae were transferred into tubes, and attached settlers were snap-frozen on petri-dishes using liquid nitrogen after excess seawater was decanted. Sample tubes and petri dishes were taken to James Cook University, (Townsville, Australia) in dry-ice, and stored at - 80◌֯C until further processing.

Frozen samples in the plates were submerged in RNAlater-ICE and allowed to thaw overnight at -20°C. Once thawed, RNAlater-ICE was removed and replaced by 2ml of *mir*Vana™homogenisation buffer. Polyps were gently scraped off the plate with a sterilized scalpel in the homogenisation buffer and transferred into a 2ml tube for immediate processing.

For both pre- and post-settlement samples, total RNA was extracted using the *mir*Vana™ miRNA Isolation Kit (Ambion) according to the manufacturer’s instructions. RNA quantity and quality were assessed using a NanoDrop ND-1000 spectrometer and denaturing gel electrophoresis using standard methods (Russell & Sambrook 2001) before being shipped on dry-ice to AGRF in Brisbane for Hi-Seq paired-end sequencing.

RNA sequencing data for all samples was first checked using a Nextflow (Di Tommaso et al. 2017) pipeline https://github.com/marine-omics/moqc. This pipeline includes fastqc (v0.11.5; https://www.bioinformatics.babraham.ac.uk/projects/fastqc/) to check raw read quality, and read-level taxonomic classification via krakenuniq (v1.0.2 ; Breitwieser, Baker and Salzberg, 2018) to determine the dominant symbiont genus and check for bacterial or eukaryotic contaminants. All samples were dominated by reads attributable to the coral host but also contained a small fraction (0.5-2%) from Symbiodiniaceae, of which *Durusdinium* was the dominant genus. No samples were flagged as having low quality base calls, excessive adapter contamination or high levels of potential contaminants (eg bacteria).

After confirming the overall quality of read data for all samples, reads were trimmed using fastp (Chen, Zhou, Chen and Gu, 2018) to remove adapters and low-quality sequences. Trimmed reads were then aligned to a combined reference database consisting of all *Acropora millepora* transcripts predicted from gene models along with the transcriptome of an unnamed *Durusdinium* species (Shoguchi et al. 2021). Alignment was performed using Bowtie2 (v2.5.0; Langmead and Salzberg, 2012) as part of RSEM (v1.3.3; Li, B., Dewey, C.N., 2011) and used to produce transcript level counts for each sample. All of these steps were run using a Nextflow pipeline https://github.com/marine-omics/morp.

Transcript counts produced by RSEM were imported into R and processed using the variance stabilising transform function in DESeq2 (vv1.38.3; Love, Huber and Anders, 2014) to produce values on a log2 scale for visualisation.

Functional annotations for *Acropora millepora* transcripts were obtained using a nextflow pipeline (https://github.com/marine-omics/moat) that is primarily based around identifying homologous proteins in Swissprot (downloaded December 2022) via blastp and blastx (NCBI Blast+ v 2.13.0). In this process, any protein with an evalue less than 1e-5 is considered a hit and the best hit is used to lookup Gene Ontology ID’s and a protein name from the Swissprot database. In addition, transcripts were annotated with conserved domains, gene family membership and an alternative Gene Ontology ID inferred using Interproscan (v5.59-91.0; Jones et al. 2014).

Principal component analysis (PCA) was conducted using the prcomp function of the stats package in R with vst normalised counts.

Weighted gene co-expression network analysis (WGCNA) was conducted with normalised data that were filtered for genes with excessive number of missing values or zero-variance using the goodSamplesGenes function of the WGCNA package (Langfelder & Horvath, 2008). The samples were clustered using the function hclust of the stats package (R Core Team, 2024) to detect outliers. One sample collected 3 hpi was removed from the analysis. To determine groups of genes +(modules) with correlated developmental gene expression profiles, a weighted gene co-expression network was constructed blockwise using the function blockwiseModules with a maximum block size of 18000 genes, a minimal module size of 18 and a mergeCutHeight set to 0.25. A signed, scale-free topological overlap matrix for each block was created by choosing a soft threshold power of 12 and a biweight midcorrelation.

To facilitate reuse and rapid exploration of the *Acropora millepora* developmental transcriptome we created an interactive web application using the shiny framework (https://shiny.posit.co/) . This application is hosted on servers provided by the Australian Research Data Commons Nectar Research Cloud to ensure that it is scalable and highly available for long-term use. Users can access the web application directly at https://amil-deview.mmb.group/ or obtain its source code and the underlying data from our github repository (https://github.com/iracooke/amil2_dev). The application is designed to facilitate access to developmental expression profiles for individual genes or groups of genes. Users can define genes of interest in two ways by either conducting a BLAST search of a sequence or by directly searching for genes with metadata matching a set of gene identifiers, Pfam domains, GO term IDs or text as part of the protein name.

Once a list of genes has been identified, the corresponding normalised gene expression data is rapidly retrieved from an sqlite relational database and used to generate an interactive display. When the number of genes is small (default <10) each gene is displayed using a separate line plot generated by the R package ggplot2 (Wickham, 2016) so that detailed changes in the expression of each gene can be carefully inspected. For larger numbers of genes (default>=10) we assume that the user is more interested in broad patterns among genes and the display changes to an interactive heatmap generated with the R package InteractiveComplexHeatmap (Gu and Hübschmann, 2022). In both cases a list of genes with annotation information is also displayed and users can select rows in this table to adjust which genes are displayed in the main display. Comprehensive data and code exporting functions are also provided so that users can easily export both data and R code to reproduce the plots seen in the app to allow further customisation by the user.

## Results

### Major transcriptional transitions visualised using DEView

During the development of *A. millepora*, the period immediately following fertilisation (0 – 5h) was characterised by rapid rounds of cell division (Figure. 1) accompanied by relatively limited changes in gene expression (Figure. 2), implying that this phase is driven largely by maternal mRNAs and proteins. After that time more extensive changes in gene expression occurred resulting in resolvable PCA clusters. After completion of gastrulation at 35hpf, embryos were spherical and became motile at 45hpf. Elongation from pear- (57hpf) to spindle-shaped larvae (69hpf) resulted in planulae that were not searching at 81hpf but were probing the environment for appropriate settlement sites at 93hpf, by which time they were competent to settle (Figure. 1). At 133hpf, settlement was induced with ethanolic CCA extract but the 133hpf transcriptome data represent the untreated state. Settlement induction caused larvae to further elongate at 15mpi before forming disc-like shapes by contracting along the oral-aboral axis. Most larvae were metamorphosed at 2hpi and by 3hpi 100% of larvae had initiated metamorphosis characterised by a six-fold symmetry around an oral pore in the centre. Whereas transcriptome data for the 0-3hpi stages clustered with those for late planula stages, data for 6 – 72hpi stages formed a discrete and well-resolved cluster (Figure. 2) during which metamorphosing polyps developed a 12-fold symmetry and formed tentacles around the oral opening. The polyp initiates calcification at the former aboral part, which becomes the calicoblastic ectoderm.

**Figure 2.**
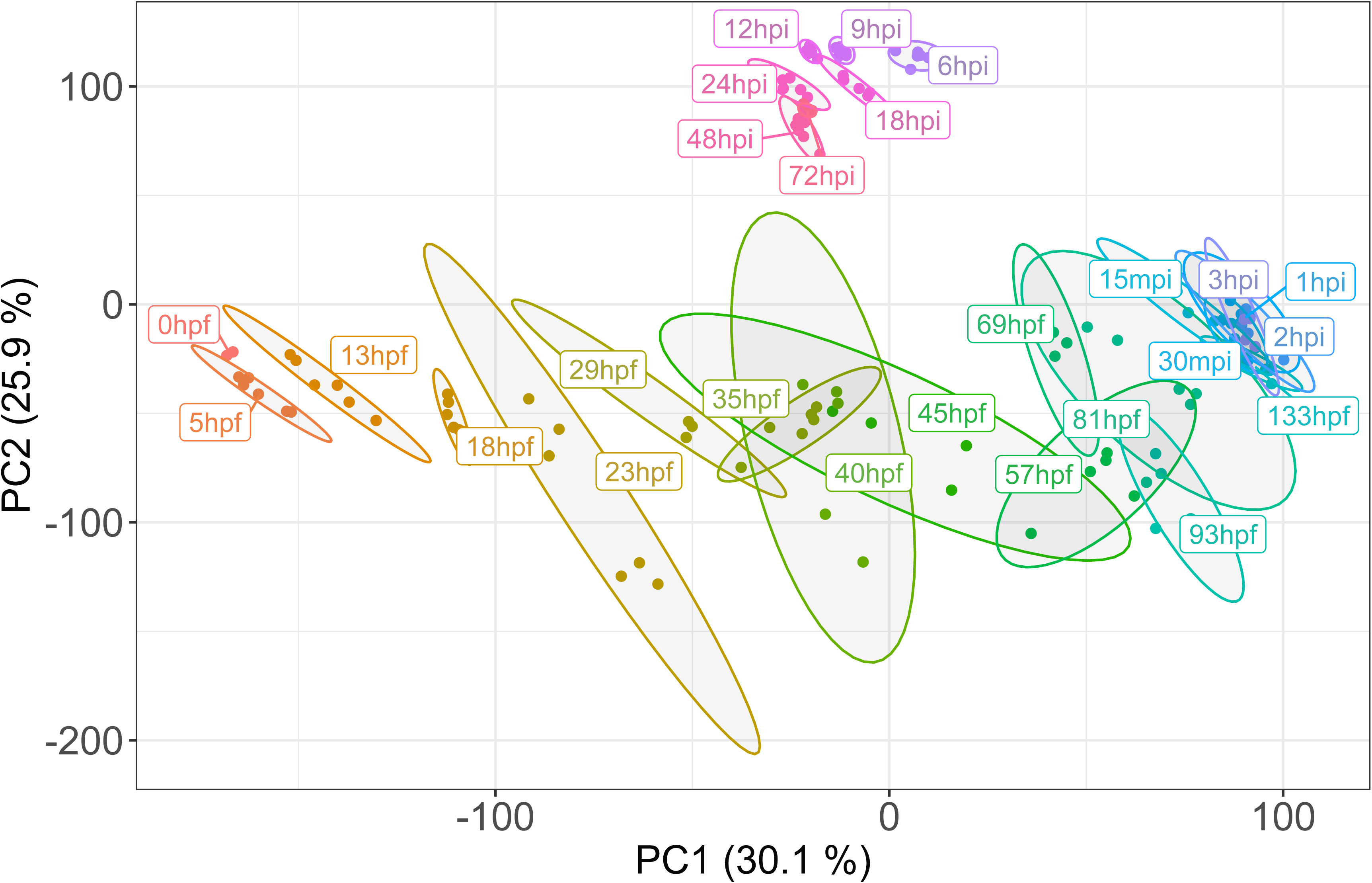
PCA analysis of the developmental transcriptomic data. Principal Components Analysis (PCA) of the developmental transcriptome data with ellipses representing 95% confidence levels. At 93 hours post fertilisation (hpf), larvae were competent to settle in response to ethanolic CCA extract and settlement was induced at the age of 133hpf. Settlement induction caused larvae to elongate at 15 minutes post induction (mpi) and 100% of larvae had initiated metamorphosis at 3 hours post induction (hpi). A major change in gene expression occurred between 3 and 6hpi.

Differential gene expression analysis (Figure 3) clearly highlights that, in addition to the expected early (13-29hpf) transcriptional events corresponding to zygotic genome activation, major transitions in gene expression also occurred at 81-93hpf and 3-6hpi. The construction of weighted gene co-expression networks (WGCNA; Langfelder & Horvath (2008), which group modules of genes according to correlation in their expression patterns, also support a 3-transition hypothesis of early coral development (Supplementary Figure 1). Whilst such analyses are informative with respect to the major transitions in development, they are less useful for exploration of the expression patterns of subsets of genes in terms of structure or function according to Pfam or Gene Ontology (GO) IDs. To facilitate these latter kinds of analyses, a software tool based on the Shiny web framework (https://shiny.posit.co/) was developed (https://amil-deview.mmb.group/).

**Figure 3.**
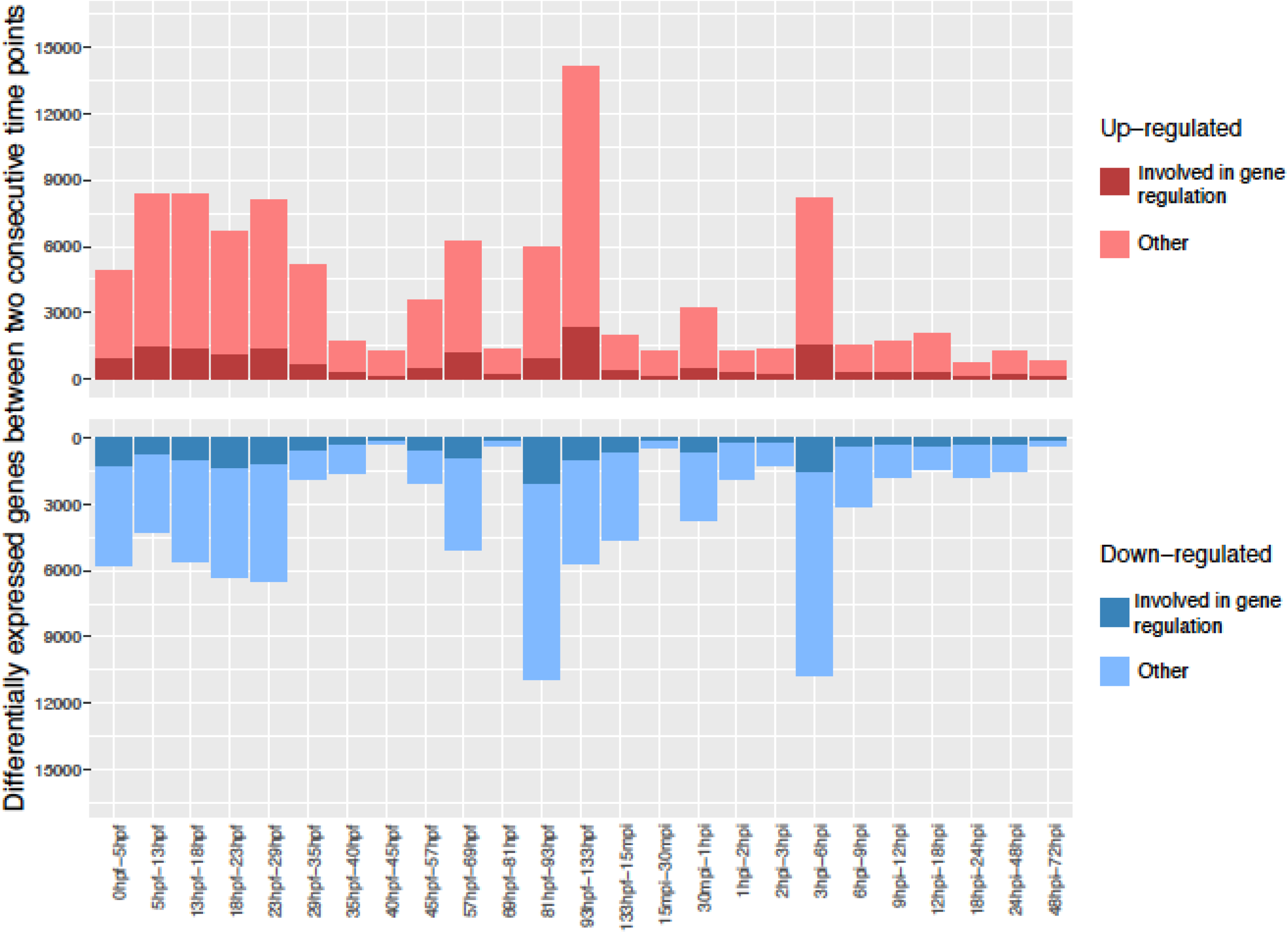
Serial up-down expression plots showing the number of genes changing in expression between consecutive time points during development. There are three periods that show major changes in expression: early development leading up to gastrulation (5-29hpf), 81-133hpf as competence to settle is developing, and 3-6hpi as polyp development is getting under way after settlement.

One powerful feature of the new software tool is that it allows the simultaneous visualisation of developmental gene expression patterns from a subset of genes defined by gene identifiers, Pfam domains or GO terms. These subsets can be selected for further analyses and comparison based on morphology, behaviour, or gene expression patterns, such as occur during gastrulation and settlement.

Whilst early zygotic gene expression accompanied gastrulation, the major transition at 81-93hpf is likely to correspond to larval competence which is the preparation for settlement and metamorphosis. Both of these transitions have previously received considerable attention (e.g. Hayward et al. 2011; Grasso et al. 2011; Strader et al. 2018). Metamorphosis was underway by 3hpi and was followed by changes in the expression of large numbers of genes (Figure 2, Figure 3). This post-settlement change in the expression patterns of many coral genes has previously received only limited attention (Ishii et al. 2022) and therefore merited closer examination. In what follows we have proceeded chronologically, with a particular focus on processes and classes of genes not discussed in previous studies.

### Maternal transcripts and the earliest transcriptional events

As in most animals, early post-fertilisation development in corals consists of rapid cell division during which the maternal cytoplasm is serially subdivided (reviewed in Ball et al. 2021), and the transcripts required to facilitate these events are largely provided maternally. Extensive zygotic gene expression appears to be initiated leading into gastrulation (i.e. around 13h post-fertilisation; Figure 1, Figure 2) corresponding to the “prawn-chip” morphology.

Rapid rounds of cell division during early development require that replication occurs quickly, generating a need for core histone proteins in large amounts to form the H2A/H2B and H3/H4 dimers that constitute the nucleosomes responsible for packaging the DNA into chromatin. As in *Hydractinia* (*Klyxum*) *echinata* (Ayers et al. 2023), histone mRNAs are amongst both the most abundant maternal transcripts and earliest expressed zygotic genes in *Acropora* (Figure 4A). Note, however, that the early zygotic (canonical) histone transcript profile is likely to be dominated by the replication-coupled types.

**Figure 4.**
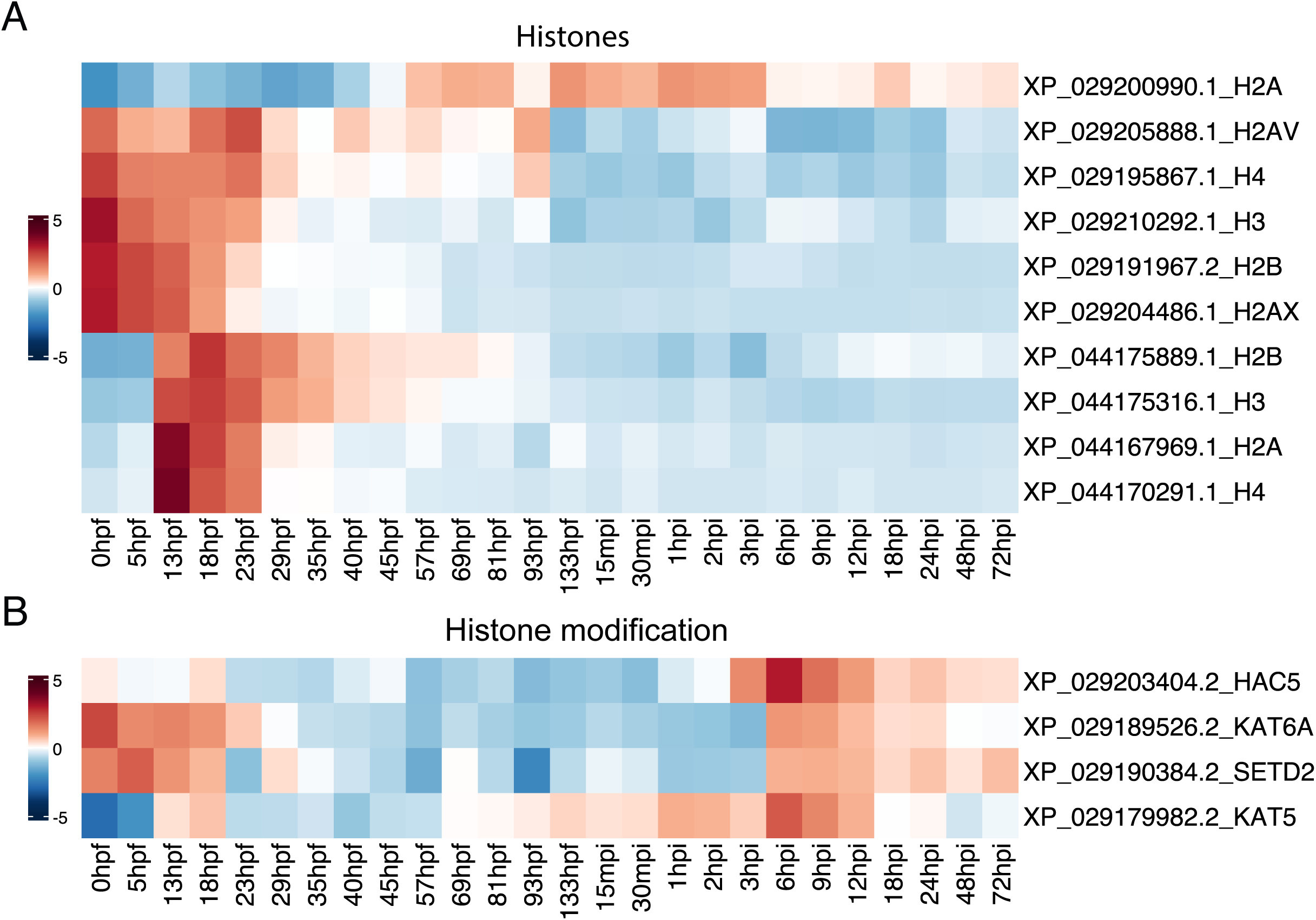
Expression of histone genes and histone modifying enzymes during the early development of *Acropora millepora*. (A) Heatmap showing expression of selected histone genes during development. mRNAs encoding canonical histones H3, H4 and H2B are present at high levels in eggs (rows 3, 4 and 5 from the top), as are H2A.X and H2A.V (but not canonical H2A) messages. Zygotic expression of the four core canonical histones begins at the start of gastrulation (i.e. the 13hpf time point) note that the set of histone genes shown in the bottom four rows are likely to represent a replication-dependent subset that may be required at high levels only early in development. Consistent with this, the top row shows data for another H2A gene, expression of which begins later in development. (B) Heatmap showing expression of histone modifying enzymes during the early development of *Acropora*. mRNAs encoding the histone acetyltransferase KAT6A the histone H3K36 methylase SETD2 are maternal (rows 2 and 3, respectively). Two other histone acetyltransferases (HAC5 and KAT5; top and bottom rows) are most highly expressed at the 6hpi time point, reflecting their possible role in activating chromatin during the post-settlement transition. Note that a second wave of expression of both KAT6A and SETD2 also occurs at that time.

**Figure 5.**
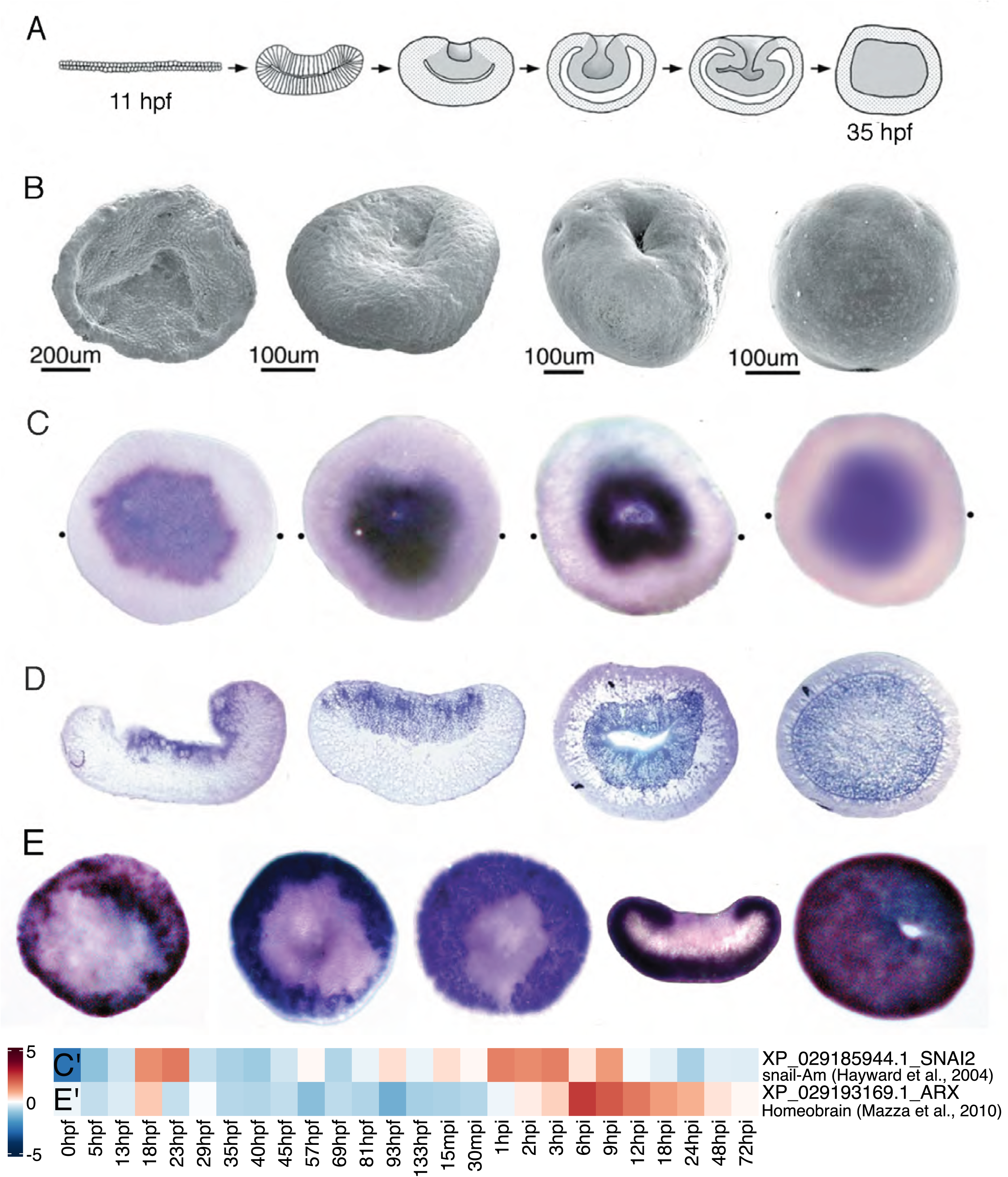
Heatmaps and in situ hybridisation provide complementary data, as illustrated by data for coral homologs of the *Drosophila* genes, snail and homeobrain. During gastrulation, these genes have complementary morphological expression patterns, *Am-snail* marking the presumptive mesendoderm (Hayward et al. 2004) and *Am-homeobrain* the presumptive ectoderm. (A) Diagrammatic vertical sections of embryos aged from early prawn chip to sphere. (B-D) Embryos aged from prawn chip to sphere (blastopore closure) viewed in three different ways. (B) Critical-point dried embryos viewed with scanning electron microscopy reveal the external morphology, (C) Whole mount in situ hybridisation reveals the spatial expression of *Am-snail*. (D) Vertical sections of in situ preparations reveal the internal distribution of the expressing tissue in the developing mesendoderm. (E) Expression of *Am-homeobrain* in the developing ectoderm revealed using whole-mount in situ hybridisation. Note that the two genes are complimentary in their expression. (C’, E’) Heatmap showing the relative expression of the two corresponding genes over the course of development. Parts of the figure were published in Hayward et al. (2004) Development Genes & Evolution 214:257-260, Springer Nature.

Transcripts encoding the core histones H2B, H3 and H4 are particularly abundant in the *Acropora* maternal mRNA complement (Figure 4A), and the zygotic expression of these histone types began early at the onset of gastrulation with a peak at 13-18hpf for H3, H4 and H2B (XP_044167100.1, XP_044175889.1) (Figure 4A). Curiously, however, canonical H2A mRNA was not maternally provisioned at significant levels, instead, one of two H2A.X homologs (H2A.X2; XP_029204486.1) and, to a lesser extent, H2A.V (H2A.F/Z; XP_029205888.1) transcripts were maternal. Levels of mRNA encoding maternal H2A.X in the unfertilised eggs were very high but fell dramatically after 13hpf as zygotic expression of canonical H2A began (e.g. XP_044167969.1), and H2A.X transcripts were essentially absent by 29hpf (Figure 4A).

Gastrulation is characterised by large scale activation of zygotic genes, requiring extensive chromatin modification. Consistent with this, high levels of mRNAs encoding homologs of the histone acetyltransferase (HAT) KAT6A (XP_029189526.2) and SETD2 (XP_029190384.2) were provided maternally (Figure 4B). The latter is the only histone methyltransferase capable of generating the H3K36me3 modification essential for transcriptional activation (Michail et al. 2025). Note that both of these genes were reactivated during the post-settlement transition discussed below.

### The MZT occurring at gastrulation marks the first major developmental transition

To date, the process of gastrulation has been the most extensively studied aspect of developmental gene expression in both corals and *Nematostella* (Technau, 2020). In situ hybridisation analyses have shown that orthologs of several key developmental regulators from bilaterians are temporally and spatially expressed in *A. millepora* consistent with conservation of function. Examples of these include *snail* (Hayward et al. 2004), *brachyury* (Hayward et al. 2015; Yasuoka et al. 2016), *forkhead* (Hayward et al. 2015) and *dpp/bmp4* (Hayward et al. 2002) (Supplementary Figure 2).

With the sole exception of *Pavona*, all complex corals examined to date go through a “prawn chip” stage of early development that corresponds to the early gastrula (Okubo et al. 2013 and references cited there; Okubo et al. 2016). At this stage, a homolog of *Drosophila snail* (XP_029185944.1) was expressed in the presumptive mesendoderm (Hayward et al. 2004). As *snail*-expressing cells are undergoing an epithelial-mesenchymal transition (EMT), one of several *Acropora* genes related to *Drosophila* aristaless (XP_044163657.1/XP_029193169.1) was expressed throughout the presumptive ectoderm (Figure 5). Hence these two genes effectively demarcate the two primary tissue layers of *Acropora* during gastrulation.

### The late planula transition corresponds to the acquisition of larval competence

During the period 81-93hpf, many genes show strong down-regulation, but this is immediately followed by the up-regulation of an even larger number of genes during the 93-133hpf time window (Figure 3).

These timepoints correspond to the acquisition of settlement competence and when *Acropora* larvae are searching for cues that identify appropriate settlement sites (eg. Tebben et al. 2015, Siboni et al. 2020, Quinlan et al. 2023, Turnlund et al. 2025) by swimming along the substratum to periodically sample it with the aboral end.

Settlement involves the aboral larval ectoderm making contact with substratum, which is likely coated with attractive and potentially hostile bacteria, and among the upregulated genes a number encode proteins with known or suspected antimicrobial properties. Among them are for example AmAMP1 (Mason et al. 2021; XP_029188806.1), the MAC-Perforin protein apextrin (Miller et al. 2007; Hayward et al. 2011; XP_044181104.1) and the quorum quenching enzyme AmNtNH1 (Mason et al. 2024; XP_029207486.1). Figure 6 summarises developmental expression data for these three genes.

**Figure 6.**
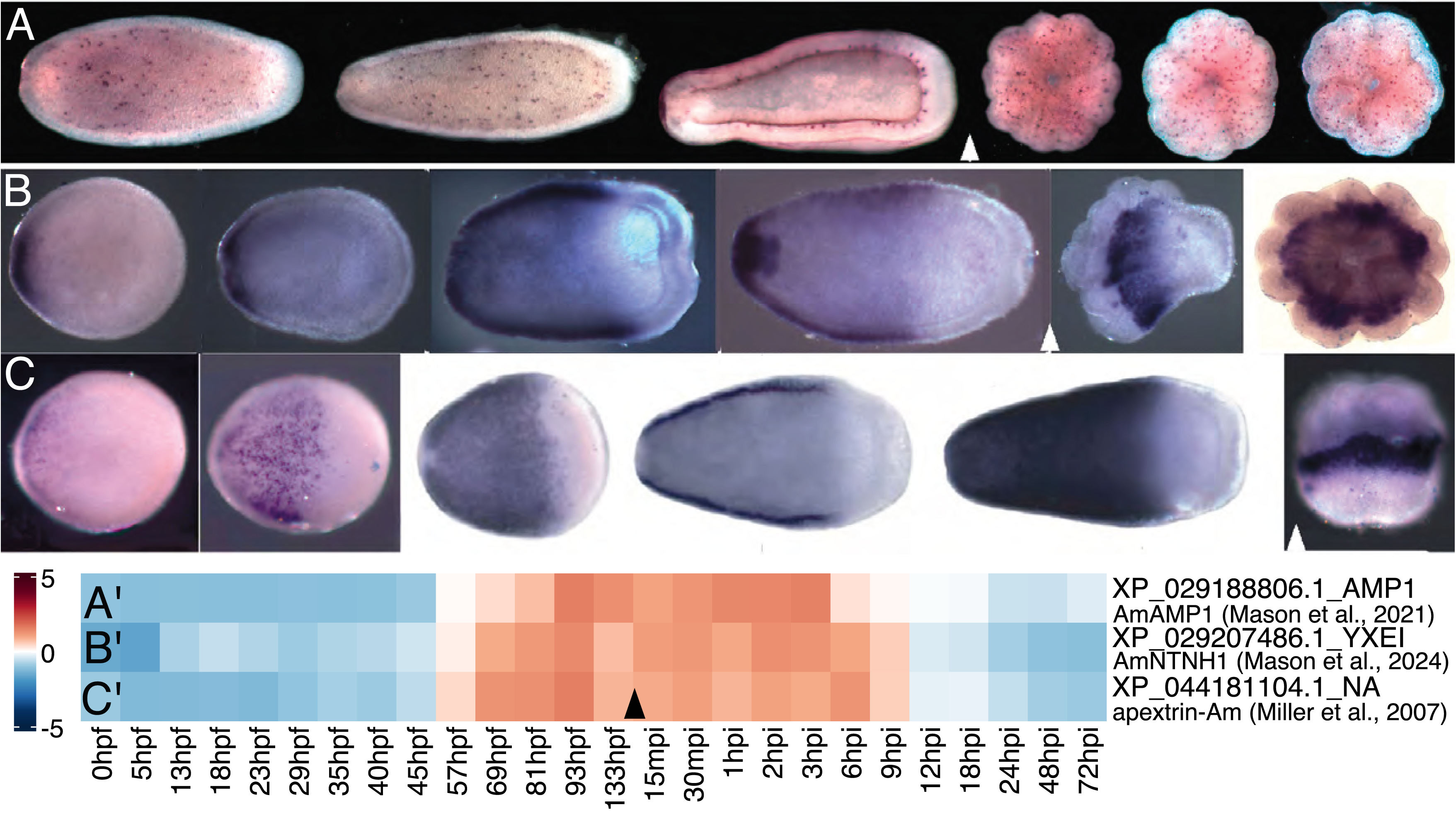
In situ hybridisation and heatmaps provide complementary data in support of roles for known and suspected anti-bacterial activities during settlement and metamorphosis in *Acropora*. The upper three rows show in situ hybridisation data for developmental stages leading into and immediately following settlement for (A) the anti-microbial protein AmAMP1 (Mason et al. (2021), (B) the quorum quenching activity AmNtNH1 (Mason et al. 2024), and (C) the MAC-Perforin protein Apextrin Miller et al. 2007; Hayward et al. 2011). A’, B’ and C’ are heatmaps showing the corresponding temporal expression levels of the three genes. The arrowheads indicate the time of settlement induction.

This study also provides the most direct evidence to date in support of Hadfield’s hypothesis (Hadfield, 2000) that coral larvae are primed for settlement and metamorphosis via prior gene expression. Whilst several thousands of genes are up or down regulated during the onset of competency (81-133hpf), settlement and the initiation of metamorphosis largely happen without changes in gene expression (Figure 3). With the caveat that datasets and technologies were much less sophisticated than those presently available, Grasso et al. (2011) came to the same conclusion for *A millepora* based on microarrays, as did Strader et al. (2018) indirectly by correlating expression data for competent larvae with settlement-induced larvae from Meyer et al. (2011).

### The post-settlement gene expression transition resembles an MZT in terms of scale and involvement of some specific components

The scale of the post-settlement transition between 3-6hpi resembles that typical of an MZT. In *Acropora*, we estimate that approximately 20% of mRNAs undergo at least a two-fold change in expression level (i.e. are either up- or down-regulated over the 3-6hpi time window). By comparison, in the zebrafish approximately 25% of maternal transcripts are cleared during the MZT (Mishima and Tomari, 2017) and the corresponding number for *Caenorhabditis* is around 33% (Baugh et al. 2003). Many of the mRNAs up-regulated over the 3-6hpi time window in *Acropora* coded for transcription factors or other regulatory proteins involved in chromatin modifications and mRNA processing.

### Chromatin modification during early development and at the post-settlement transcriptional transition

Major changes in gene expression patterns also generally involve chromatin modification events, so histone modification and DNA methylation changes are to be expected. Homologs of a number of key chromatin modifiers are differentially expressed at the 3-6hpi transition.

In many animals, homologs of TET (ten-eleven translocation) are the primary agents responsible for demethylation of 5-methylcytosine residues (MeCpG) in DNA and are required for gene activation (Mohr et al. 2011; Wu and Zhang 2017; Jessop et al. 2018). As well as a transient earlier (5hpf) window of expression leading into the early MZT, the *Acropora TET* homolog (XP_029179484.2) was strongly upregulated at the 6hpi time point (Figure 7A), suggesting a role for the *Acropora* TET homolog in relieving repression by erasing pre-existing MeCpG marks.

**Figure 7.**
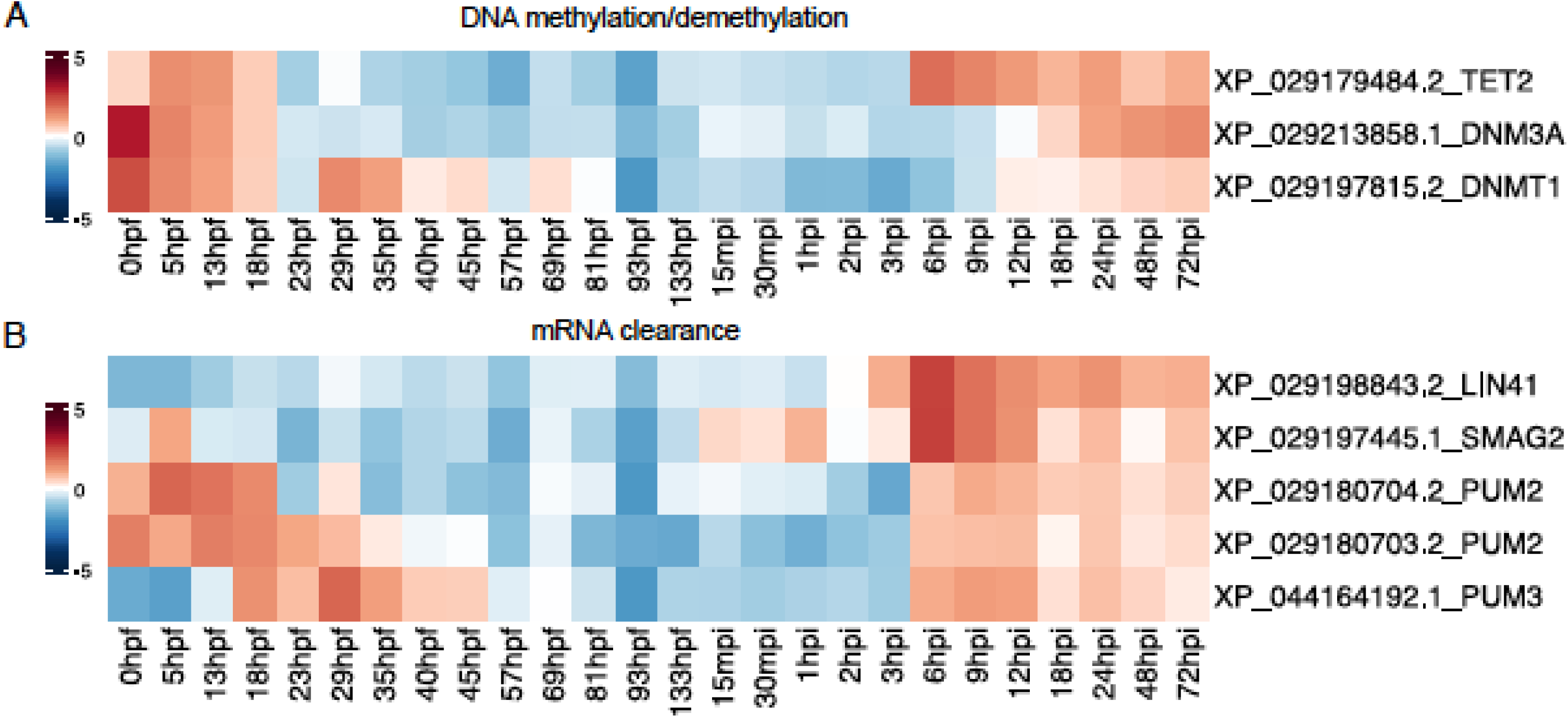
Heatmaps for genes implicated in (A) DNA methylation /demethylation and (B) clearance of mRNAs and proteins are consistent with major changes in gene activity occurring at 3-6hpi. The upper panel (A) illustrates relative expression levels of the *Acropora* homologs of ten-eleven-translocase (tet), the primary role of which is removal of CpG methylation marks, the de novo DNA methyltransferase, DNMT3, and the maintenance DNA methyltransferase, DNMT1. Although all three activities are expressed during gastrulation, TET is most heavily expressed at the 6hpi time point. The lower panel (B) shows relative expression levels of *Acropora* homologs of the conserved mRNA clearance activities, Smaug and Pumilio, and the TRIM-NHL ubiquitin ligase, TRIM71/LIN41. Smaug and TRIM71/LIN41are most heavily expressed at the 6hpi time point; three gene predictions annotated as Pumilio homologs (corresponding to at least two distinct gene products) were also up-regulated at that time.

DNA methylation is generally associated with repression of transcription, the enzymes associated with maintenance and *de novo* DNA methylation being the DNA methyltransferases DNMT1 and DNMT3, respectively. In the case of *Acropora*, mRNAs encoding the DNA methylases DNMT1 (XP_029197815.2) and DNMT3a (XP_029213858.1) were maternally provided, zygotic expression of DNMT1 being detected between 29hpf and 12hpi, despite having faded by 81hpf. Both DNMTs were upregulated 12 to 18hpi (Figure 7A) expression of DNMT3a appeared to be stronger than that of DNMT1 (Figure.7A)

### mRNA degradation during major transitions in development

Perhaps the major characteristic of MZTs is the elimination of a subset of maternally provided mRNAs (often a major proportion) to facilitate the transition to the zygotic developmental program. The mRNA elimination process employs a range of mechanisms to destabilise or degrade transcripts, but some conserved players have been identified during major transitions, including homologs of the *Drosophila* genes *Pumilio* (Gerber et al. 2006; Rabani et al. 2017) and *Smaug* (Chen et al. 2014) as well as the *Caenorhabditis* protein TRIM71/LIN41 (Spike et al. 2022).

Transcripts annotated as *Pumilio* (XP_029180703.2 and XP_029180704.2) were maternally provided but were essentially cleared after 29hpf, whereas a distinct Pumilio homolog (XP_044164192.1) was not represented in the maternal mRNA complement but was expressed from 18hpf (Figure 7B). Each of these Pumilio homologs was up-regulated over the 3-6hpi time window after earlier periods of activity.

Members of the Smaug/SAMD4 family of proteins are highly conserved (yeast to man) regulators of mRNA stability and transcriptional repressors that act in development by binding to specific RNA sequence motifs. Smaug proteins have been implicated in erasing maternal transcripts during the MZT in the early development of diverse animals (Chen et al. 2014; Chille et al. 2021) implying that the Smaug homolog fulfills a similar function in the early development (around 4hpf) of the coral *Montipora capitata*. Although an early “flutter” of Smaug expression, possibly corresponding to that reported in *Montipora* (Chille et al. 2021), was observed in *Acropora* at 5hpf, Smaug (XP_029197445.1) was predominantly expressed at 15mpi - 6hpi (peak) (Figure 7B). A second transcript annotated as Smaug (XP_044163536.1) was mildly up-regulated at 13-18hpf.

Members of the TRIM71/LIN41 family of E-3 ubiquitin ligases are highly conserved RNA-binding proteins (Aviv et al. 2003) that facilitate major developmental transitions in *Caenorhabditis* (Spike et al. 2022). At least one member of this gene family (XP_029198843.2/XP_029198844.2) is up-regulated at the 3-6hpi time point during *Acropora* development, suggesting that they may also be involved in protein clearance at the 3-6hpi transition.

### Small RNA processing pathway components

Small RNAs (miRNAs, siRNAs and piRNAs) have important regulatory functions during the development of many bilaterians (Bartel, 2004) and this is likely to also be the case in *Acropora* given that at least some components of the miRNA processing pathway are known to be differentially expressed during the development of *Nematostella* (Praher et al. 2017). In cnidarians, the biogenesis of miRNAs follows a plant-like pathway (Moran et al. 2017), but homologs of some of the enzymes in the animal miRNA processing system are also present and may also participate in miRNA synthesis.

To address the possibility of miRNAs participating in gene regulation during early coral development, the expression of components of both the plant and animal processing pathways was investigated (Figure. 8).

**Figure 8.**
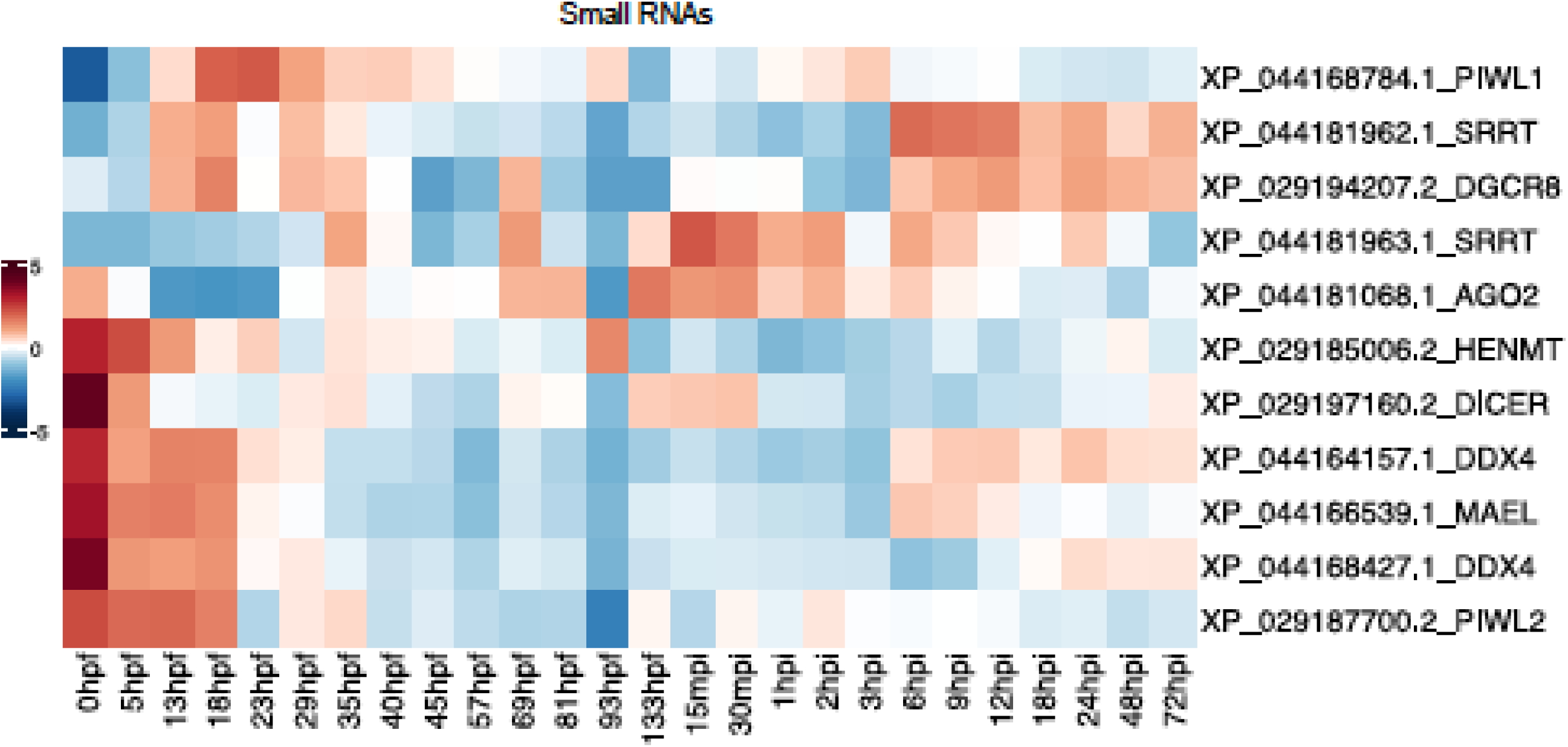
Expression of genes involved in small RNA (miRNA and piRNA) processing during early development. mRNAs encoding HEN1 and Dicer homologs (rows 6 and 7) were present at high levels in eggs, as were those encoding homologs of Maelstrom, Vasa/DDX4.1 and .2, and Piwi2, which function primarily in piRNA-mediated transposon silencing. Other miRNA processing activities, including homologs of Serrate/SRRT and DGCR8/Pasha, were also expressed in embryos, larvae and during post settlement life, implying extensive involvement of miRNAs in regulating gene activities during early coral development.

Some of the small RNA processing pathway components were provided maternally, levels of transcripts annotated as Dicer-like (XP_029197160.2) and the HEN1 methyltransferase (HEN1MT; XP_029185006.2) being present at high levels. In cnidarians, HEN1 is responsible for methylation of the 3’-ends of both miRNAs and piRNAs, resulting in their stabilisation. The observed upregulation of HEN1 at 93hpf (Figure. 8) is consistent with a role in metamorphosis of *Acropora* larvae, as in *Nematostella* (Modepalli et al. 2018).

The maternal mRNA repertoire also included an Argonaute homolog (XP_044181068.1) and a *Acropora* Piwi-like protein (XP_029187700.2/ XP_029187701.2) (Figure. 8) that likely function together in transposon silencing during early development, as in *Nematostella* (Fridrich et al. 2020; Bamberger et al. 2022). The high abundance of mRNA encoding a homolog of the Maelstrom protein (XP_044166539.1) also attests to the importance of the piRNA-mediated transposon silencing pathway (Matsumoto et al. 2015; Sato and Siomi, 2015) during the earliest stages of coral development.

Although not maternal, other miRNA pathway components were expressed very early in development (Figure 8). Transcripts annotated as DGCR8/Pasha (XP_029194207.2) and SRRT/Serrate (XP_044181962.1) were first detected as the early embryo was going into gastrulation at about 13hpf. For both of these genes, other transcripts carrying the same annotation appeared to be expressed during different stages of development. In the case of SRRT/Serrate, XP_044181962.1 is expressed in the early gastrula but most heavily after the initiation of metamorphosis (6hpi). A second isoform/splice variant (XP_044181963.1) was most highly expressed very early in the settlement process (15mpi - 2hpi) – i.e. prior to what Ishii et al. (2022) referred to as the “point of no return”.

Similarly, the DGCR8/Pasha paralogs or splice variants XP_029194207.2 and XP_029194208.2 seem to have distinct expression profiles during development (data not shown).

In addition to the maternal Piwi-like mRNA (XP_029187700.2/ XP_029187701.2), a second (non-maternal) Piwi (XP_044168784.1/ XP_044168785.1/ XP_044168786.1) was expressed from the early gastrula stage, peaking in expression as the maternal type was fading.

## Discussion

The study of early coral (*A. millepora)* development presented here was based on gene expression profiles obtained for 26 developmental stages spanning the period between unfertilised eggs and 3-day-old primary polyps. Downstream analyses via a DEView app revealed three major transcriptomic transitions - the maternal-zygotic transition, the acquisition of competence by planulae (93-133hpf) and a post-settlement transition (3-6hpi) that has not previously received significant attention. This latter novel transition in gene expression resembled an MZT in terms of scale and the involvement of some key genes. In addition to regulatory genes involved in chromatin modification and RNA processing, we focused here on genes likely to have protective roles, specifically those potentially involved in photoprotection, anti-microbial activity and transposon silencing.

In many animals, mRNAs encoding canonical histones (including H2A) as well as the corresponding proteins are provided maternally at sufficient levels to support the earliest cell division events. To our knowledge, finding that in *Acropora*, an mRNA encoding one H2A.X variant (H2A.XII) replaced that for canonical H2A in the eggs is novel. H2A.X is a histone variant whose best-defined role is in DNA repair (reviewed in Lorkovic and Berger, 2017; Herchenröther et al. 2023). It is closely related to canonical H2A but distinguished from it by the presence of a characteristic C-terminal sequence extension that includes a serine residue subject to phosphorylation generating the gammaH2A.X form which is required for binding at double-stranded DNA breaks (Osakabe and Molaro, 2023). The enzymes responsible for this phosphorylation of H2A.X are Phosphoinositide-3-Kinase-related protein kinases (PIKKs), and at least one PIKK-related protein kinase (XP_029200091.2) is provided maternally in *Acropora*, though it is unclear whether H2A.X is a substrate for this activity.

Whilst most coral spawning occurs during the night, the gamete bundles (or eggs) released are buoyant so that the earliest developmental events occur near the water’s surface. In the present study, *Acropora* larvae became motile 40-45hpf by which time they would have been exposed to two days of direct sunlight. Early developmental stages of corals therefore face exposure to UV radiation at levels that can cause DNA damage, and H2A.X2 mRNA may be maternally provided at high levels to facilitate maintenance of DNA integrity through the earliest developmental stages. Chille et al. (2021) noted high representation of GO terms associated with DNA repair in the maternal mRNA complement of *Montipora* but did not identify specific genes/proteins. GFP-related proteins are also thought to be involved in DNA protection in corals (Salih et al. 2000; Palmer et al. 2009; Bou-Abdallah et al. 2006; Clarke et al. 2024) and, consistent with the study of Strader et al. (2018), at least four distinct types are expressed during the larval and early polyp stages. Although mRNAs encoding GFPs did not appear to be maternally provided at significant levels, the pigmentation of the eggs of many coral species is consistent with the idea of maternal provision of photoprotective agents that likely include GFP-related proteins, which are known to be long-lived.

Note that in *Acropora* there is a second H2A.X-like gene (H2A.X1; XP_029200990.1 - annotated as H2A-like) that is expressed only at relatively low levels during early development. This second H2A.X isoform (XP_029200990.1) has an N-terminal tyrosine residue replacing the phenylalanine in XP_029204486.1 and has close homologs in *Hydra* (CDG71330.1; 86.47% protein identity to *Acropora*) and *Hydractinia* (now *Klyxum*; AOF43153.1; 87.02% identity) and, like those, is presumed to be the main isoform expressed during post-larval life. In *Hydractinia* (now *Klyxum*) *echinata*, the H2A.X variant AOF43152.1 is expressed specifically in female germ cells along with canonical histone H2A (Torok et al. 2016) and may be functionally equivalent to the maternally provided *Acropora* H2A.X homolog XP_029204486.1 (protein sequence identity of 68.15%).

### Comparisons with published studies of coral development

The most interesting observation made here was the extent and nature of the post-settlement transcriptional transition observed at 3-6hpi. This major change in gene expression is more apparent than in previous studies which either did not include post-settlement stages (Strader et al. 2018; Chille et al. 2021) or sampled sparsely pre-settlement (Huffmeyer et al. 2025) or after settlement induction (Ishii et al. 2022, Meyer et al. 2011) (Table 1). Furthermore, comparisons between the work presented here and previously published papers in this field (Table 1) are complicated due to the diversity of species studied, differing temperatures during development, differing sampling points and differing inducers used to trigger settlement and metamorphosis.

Development has been studied primarily in members of the three coral genera *Acropora, Montipora* and *Porites*. Whilst these are all members of the “superfamily” Complexa, and the first two belong to the same coral family (Acroporidae), both *Montipora* and *Porites* spp. maternally transmit Symbiodiniaceae whereas *Acropora* spp. acquire their symbionts from the environment during early life.

Chille et al. (2021) focussed on MZT and therefore studied gene expression changes during development of *Montipora capitata* from unfertilised egg to planula in addition to adults. Some of their MZT results are consistent with those presented here for *Acropora*, for example the expression of Smaug (Figure 7B) and Brachyury (Supplementary Figure 2) in prawn chip and early gastrula. However, the rate of development of *M. capitata* observed was apparently slower than that of *Acropora*. *Montipora capitata* reached the late gastrula stage after 1.5 days, whereas *Acropora* generally reaches the corresponding stage earlier (around 30hpf). This demonstrates the complication of relating developmental stages across datasets where interspecific differences can be compounded by the presence/absence of maternal photosymbionts and different rearing temperatures, as higher temperatures accelerate development. Using GO-term data, Huffmeyer et al. (2025) recognised three distinct (but overlapping) phases of gene expression during the early development of *Montipora*.

Since morphology reflects changes in gene expression, we have found that comparisons of gene expression based on morphology are more meaningful than those based on time, even in the single species *A millepora*. The post-settlement transition remained unexplored in the study of Chille et al. (2021) since the focus on MZT did not require the sampling of post-settlement stages. The 3-6hpi gene expression transition observed here may correspond to what Ishii et al. (2022) refer to in their study as “the point of no return” (PNR), a point in post-settlement development after which reversibility by aborting settlement and returning to the motile phase is lost. Whilst the Ishii et al. (2022) paper focussed on gene expression changes between three time points during metamorphosis in *Acropora*, comparison between their results and those presented here is complicated by the differing triggers used to drive the process. For *Acropora* spp., external exposure of competent larvae to the *Hydra* GLWamide (Hym-248), directly triggers metamorphosis, essentially overriding the need for external settlement cues (Iwao et al. 2002; Grasso et al. 2011; Attenborough et al. 2019). Whilst Ishii et al. (2022) induced metamorphosis using Hym-248, the study described here triggered settlement by addition of CCA extract, which induces searching behaviour rather than directly driving metamorphosis (Grasso et al. 2011). Hence it is not possible to directly compare developmental timing between these two studies. The 1hpi and 6hpi time points in the Ishii paper are likely to correspond to later time points in the present study, as the duration of the settlement period must be several hours longer when using CCA extract compared to Hym-248. The PNR inferred by Ishii as occurring at 4hpi may correspond to the 3-6h transition identified in the present work but note that rates of larval development can be strongly influenced by temperature. The differences in gene expression between untreated (0 hr) and 1 hour after induction described in the Ishii et al. paper are therefore likely to encompass a number of distinct time points in our study, so subtle transcriptional changes occurring during the searching and very early settlement phases were presumably not noted by Ishii et al. (2022).

### The post-settlement transition – a cnidarian equivalent of the *Caenorhabditis* “juvenile to adult” transition?

In *Acropora*, the transition from larvae to recruits at 3-6hpi has several parallels to maternal-zygotic transitions (Vastenhouw et al. 2019) regarding the removal of earlier transcripts and genome activation appropriate for stages following the transition. A precedent for such a later transition is provided by *Caenorhabditis elegans*, where it is recognised as the juvenile to adult transition (Spike et al. 2022).

Above, the involvement of homologs of both *Drosophila* Smaug and Pumilio in the mRNA clearance processes during MZT and the juvenile to adult transition at 3-6hpi were highlighted. In addition, two transcripts with the same sequence (XP_029198843.2 and XP_029198844.2) annotated as a member of the TRIM71/LIN41 family of RNA-binding proteins were upregulated during the 3-6hpi transition. Other members of this protein family have important roles in the regulation of both oocyte to embryo (Spike et al. 2022) and juvenile to adult (Aeschimann et al. 2019) developmental transitions in *Caenorhabditis*. As in *Caenorhabditis* (Spike et al. 2022), this *Acropora* TRIM71/LIN41-related protein is likely to be involved in regulatory changes at the 3-6hpi transition, and on this basis deserves further exploration.

In addition, peaks in the expression of the DNA demethylase TET (Figure 7A) and of SETD2 (XP_029190384.2) as well as homologs of the histone acetyltransferases HAC5 (XP_029203404.2), KAT5 (XP_029179982.2) and KAT6A (XP_029189526.2) (Figure 4B) around 3-6hpi imply extensive chromatin modification at that time. Presumably, the corresponding proteins function in facilitating transcription of genes required for early polyp development (e.g. He et al. 2024). Transcription factors (TF) strongly up-regulated at the 3-6hpi time point include genes annotated as homologs of FoxB1 (XP029196672.1) and FoxO (XP_029208531.2), Aristaless (XP_029193170.1), Rax (XP_029193171.1), Meis2 (XP_029187327.1) and Otx (XP_029190339.1), whilst homologs of Emx (XP_029211983.2), Pitx2 (XP_029179676.1) and Brn3C (XP_029213135.1) were simultaneously down-regulated.

### Limitations: Heatmaps are useful but tell only part of the story

A caveat in making inferences from heatmaps across developmental stages, and more generally from whole-larvae overall expression levels, is that important subtle changes may be missed. Particularly with transcription factors or other regulatory molecules, low overall levels of expression may have major effects, especially when restricted to subsets of cells. Consequently, interpreting developmental gene expression requires not only temporal data, but also spatial information from the application of in situ hybridisation or other techniques. This is clearly illustrated in Supplementary Figure 2, which provides expression data for genes implicated in conserved functions during gastrulation.

Whilst in situ hybridisation data show expression of these four genes in smaller subsets of genes across all developmental stages between gastrulation and metamorphosed recruit, the heatmaps show that the expression maxima occur post-settlement rather than at gastrulation around 13 – 18hpf. Only a weaker peak in expression of Brachyury at 13hpf corresponds to the early gastrulation window of expression visualised via in situ hybridisation (Hayward et al. 2015). Consequently, visualisations of spatial expression via in situ hybridisation are required to study the function of selected genes. For example, whilst gene expression data presented in the heatmap suggested functions post-settlement, ISH visualisations revealed that temporal expression of an *Acropora* arx-related gene (XP_044163657.1/XP_029193169.1) during gastrulation closely complements that of snail (Figure 5). This *Acropora* sequence is referred to here as homeobrain, based on its orthology with *Nematostella* XP_048583705.1 (Mazza et al. 2010). Snail and homeobrain respectively demarcate the early mesendoderm and presumptive ectoderm during gastrulation (Figure 5), but both are also expressed at higher levels after settlement-induction where snail expression reaches its expression peak earlier than in the case of homeobrain. Assuming these two genes fulfill similar function during gastrulation and post-settlement, this suggests that a cell migration such as in an epithelial-mesenchymal transition occurs during the initiation of metamorphosis. The upper limit of expression in a heatmap is determined by the genes included in that map so important changes in gene expression may not be as apparent as if the gene was plotted against itself.

### Expression of the small RNA processing machinery during early development

Despite deep evolutionary divergence between the sea anemones (Actiniaria) and the coral/corallimorpharian lineage, many of the molecular processes underlying early development in *Acropora* closely parallel those described in *Nematostella* (Hayward et al. 2015), and this appears to extend to the machinery for processing small RNAs (miRNAs and piRNAs). However, some differences were observed in the timing of expression of several of genes implicated in small RNA processing.

Whilst mRNAs for Vasa1, Maelstrom and Piwi are provided maternally in *Acropora* and *Nematostella*, Vasa2 (aka Vasa-like or DDX4) is also maternal in the coral whereas the corresponding mRNA was not detected until around gastrulation in *Nematostella* (Praher et al. 2017). Whether Dicer/DCL is maternal is unclear in the case of *Nematostella*, whereas mRNA of the *Acropora* homolog of *Nematostella* Dicer 2 (AGW15596.1), encoding XP_029197160.2, is abundant in *Acropora* eggs. In summary, the preliminary data presented here indicate that all or most of the small RNA processing machinery is either maternal or expressed very early in *Acropora* development. It is therefore likely that, in addition to roles for some of these components in transposon silencing, miRNAs are likely to have important regulatory roles in the early development of *Acropora*.

### Heatmaps enable identification of candidate genes for key roles in early development

A major strength of the DEView app is that it provides a means of scanning classes of genes individually or collectively for their expression patterns across early coral development, allowing identification of candidates for roles in specific processes, including settlement induction and the initiation of calcification. Whilst significant progress has recently been made, we have as yet a relatively limited understanding of the nature and recognition of settlement cues (Hadfield, 2011; Tran and Hadfield, 2012; Ball et al. 2021; Turnlund et al. 2025). The larvae of many corals select appropriate settlement sites by detection of small molecules that either favour or discourage settlement (e.g. Morse et al. 1988; Negri et al. 2001; Kitamura et al. 2007; Birrell et al. 2008; Kitamura et al. 2009; Tebben et al. 2015, Quinlan et al. 2023). Most likely the outcome reflects a combination of molecules acting as ligands for specific receptors that are expressed in competent larvae and downregulated after settlement and metamorphosis. Receptors that potentially could be involved in settlement cue reception and downstream signalling pathways, respectively, are G-Protein coupled receptors (GPCRs) and nuclear receptors. With the help of the DEView app, search terms such as “G-protein coupled receptor” or specific Pfam domains can be used as filters to search the developmental transcriptome to identify genes that were upregulated at competency (i.e. 93hpf) consistent with their potential involvement in settlement cue perception. Searching the *Acropora* data using the terms PF00104 Ligand-binding domain of nuclear hormone receptor and PF00105 Zinc finger, C4 type (two domains) yielded 21 hits corresponding to 14 loci (Supplementary Figure 3). Nuclear receptors play key roles in metamorphosis in both insects and some vertebrates, potentially acting in the internal transmission of metamorphosis signals rather than their external reception. Of the 14, seven were upregulated at 133hpf and all of these mRNAs appear to be cleared at the 3-6hpi transition so could have roles in the cue recognition pathway and/ or metamorphosis.

The *Acropora* developmental transcriptome data, together with the DEView app, have provided novel insights into early coral development and allowed identification of candidate genes for key roles. We hope that the availability of this app via the web will lead to more extensive exploration of the data and ultimately facilitate much better understanding of the molecular processes underlying the development of the coral body plan.

For a range of organisms, including some corals, considerably more advanced genomic and transcriptomic resources are now available than could have been imagined even a few years ago. However, despite the developmental transcriptomic data provided here being to our knowledge the most comprehensive available for any coral, users need to be aware that the quality of the *Acropora millepora* dataset falls short of those available for the model invertebrates *Drosophila* and *Caenorhabditis*. Nevertheless, the developmental novelties uncovered here provide clear indications of how useful the dataset can be in facilitating coral developmental research.

## Acknowledgements

The authors gratefully acknowledge the support of the Australian Research Council under Discovery Grant DP170104734, SuperScience Grant (round 2) FS110200046 and Centres of Excellence Grant CE14100020. We also thank Gergely Torda and the staff of the Orpheus Island Research Centre for fieldwork assistance. Some of the in situ images presented here were produced by Lauretta Grasso and Nikki Hislop.

## Availability of data and materials

### Data accessibility statement

To facilitate use and rapid exploration of the *Acropora millepora* developmental transcriptome we created an interactive web application that is hosted on servers provided by the Australian Research Data Commons Nectar Research Cloud to ensure that it is scalable and highly available for long-term use. Users can access the web application directly at https://amil-deview.mmb.group or obtain its source code and the underlying data from our github repository (https://github.com/iracooke/amil2_dev).

## Author Contributions

Designed research: AM, DH, BL, EB, DM

Carried out field work: AM, DH, BL, EB

Laboratory work: AM, BL

Performed bioinformatic analyses: SF, AM, DH, IC

Contributed new analytical tools: IC

Analysed the data: AM, RB, MG, DH, DM

Wrote the paper: DH, RB, EB, IC, DM (lead)

---

**Supplementary Figure 1.** Weighted gene co-expression network analysis (WGCNA) clustered the developmental gene expression data into 39 modules based on correlated expression patterns. The number of genes in each module range from 18 (plum module) to 5851 (turquoise module) and the grey module contains all genes that did not fit into any other module. The WGCNA heatmap indicated that there is a major change in gene expression between 3 and 6 hours post induction (hpi) and also showed that settlement induction at the age of 133 hours post fertilisation (hpf) did not trigger an immediate strong change in gene expression 15 minutes post induction (mpi).

**Supplementary Figure 2.** Genes previously characterised in the context of gastrulation illustrate the need for spatial as well as temporal expression data. In situ data for the *Acropora* homologs of several genes imply conservation of function during gastrulation with Bilateria. (A) BMP2/4 (BMP2B), (B) Forkhead (FOXA2), (C) Goosecoid (GSC) and (D) Brachyury (TBXT). However, heatmaps show that, for most of these genes, rather than at gastrulation (around 13 – 18 hpf) the expression maxima occur post-settlement. Only one of the genes characterised in the context of gastrulation (snail) has a major peak of expression at the late “prawn chip” (gastrula) stage (Figure 5) – which is consistent with in situ data (Hayward et al. 2004) suggesting expression throughout the presumptive mesendoderm. The in situs were previously published in Hayward et al. (2015) Developmental Biology 399:337-347 (Elsevier).

**Supplementary Figure 3.** Differential expression of genes encoding nuclear receptors during coral development. Searching the *Acropora* data using the terms PF00104 Ligand-binding domain of nuclear hormone receptor and PF00105 Zinc finger, C4 type (two domains) yielded hits corresponding to 14 loci. Two of these loci code for predicted protein isoforms with distinct expression profiles (XP_029208454.1 and XP_029208455.1 are protein isoforms from the locus LOC114972094; XP_044184859.1 and XP_044184860.1 are isoforms from the locus LOC114969161). Consistent with possible roles in cue recognition and/ or metamorphosis, seven of the 14 genes were upregulated at 133hpf. All of these mRNAs were cleared at the 3-6hpi transition.

## References

Aeschimann F, Neagu A, Rausch M, Großhans H. *let-7* coordinates the transition to adulthood through a single primary and four secondary targets. Life Sci Alliance. 2019; 2(2):e201900335.

Attenborough RMF, Hayward DC, Wiedemann U, Foret S, Miller DJ, Ball EE. Expression of the neuropeptides RFamide and LWamide during development of the coral Acropora millepora in relation to settlement and metamorphosis. Dev Biol. 2019; 446: 56–67.

Aviv T, Lin Z, Lau S, Rendl LM, Sicheri F, Smibert CA. The RNA-binding SAM domain of Smaug defines a new family of post-transcriptional regulators. Nat Struct Biol. 2003;10(8):614–21.

Ayers TN, Nicotra ML, Lee MT. Parallels and contrasts between the cnidarian and bilaterian maternal-to-zygotic transition are revealed in Hydractinia embryos. PLoS Genet. 2023;19(7):e1010845.

Ball EE, Hayward DC, Bridge TCL, Miller DJ. *Acropora* – the most studied coral genus. In: Boutet A, Schierwater B (eds). Handbook of Marine Model Organisms in Experimental Biology - Established and Emerging. 2021. Boca Raton (FL): CRC Press. p. 173–193. DOI: 10.1201/9781003217503-10.

Ball E, Hayward D, Reece-Hoyes J, Hislop N, Samuel G, Saint R, Harrison P and Miller, D. Coral development: from classical embryology to molecular control. Int J Dev Biol. 2002;46:671–678.

Ball EE, Hayward DC, Saint R, Miller, DJ. A simple plan—cnidarians and the origins of developmental mechanisms. Nat Rev Genetics. 2004;5(8):567–577.

Bamberger C, Pankow S, Yates JR 3rd. Nvp63 and nvPIWIL1 suppress retrotransposon activation in the sea anemone Nematostella vectensis. J Proteome Res. 2022; 21(11):2586–2595.

Bartel DP. MicroRNAs: genomics, biogenesis, mechanism, and function. Cell. 2022; 116(2):281–97.

Baugh LR, Hill AA, Slonim DK, Brown EL, Hunter CP. Composition and dynamics of the Caenorhabditis elegans early embryonic transcriptome. Development. 2003; 130(5):889–900.

Birrell CL, Mccook LJ, Willis BL, Harrington L. Chemical effects of macroalgae on larval settlement of the broadcast spawning coral Acropora millepora. Mar Ecol Prog Ser. 2008;362:129–137.

Bou-Abdullah F, Chasteen ND, Lesser MP. Quenching of superoxide radicals by green fluorescent protein. Biochim Biophys Acta. 2006;1760(11):1690–5.

Breitwieser FP, Baker DN, Salzberg SL. KrakenUniq: confident and fast metagenomics classification using unique k-mer counts. Genome Biol. 2018;19:198.

Chen L, Dumelie JG, Li X, Cheng MH, Yang Z, Laver JD, Siddiqui NU, Westwood JT, Morris Q, Lipshitz HD, Smibert CA. Global regulation of mRNA translation and stability in the early Drosophila embryo by the Smaug RNA-binding protein. Genome Biol. 2014;15(1):R4.

Chen S, Zhou Y, Chen Y, Gu J. fastp: an ultra-fast all-in-one FASTQ preprocessor. Bioinformatics. 2018;34:i884–i890.

Chille E, Strand E, Neder M, Schmidt V, Sherman M, Mass T, Putnam H. Developmental series of gene expression clarifies maternal mRNA provisioning and maternal-to-zygotic transition in a reef-building coral. BMC Genomics. 2021; 22(1):815.

Clarke DN, Rose NH, De Meulenaere E, Rosental B, Pearse JS, Pearse VB, Deheyn DD Fluorescent proteins generate a genetic color polymorphism and counteract oxidative stress in intertidal sea anemones. Proc Natl Acad Sci USA. 2024; 121(11):e2317017121.

Di Tommaso P, Chatzou M, Floden EW, Barja PP, Palumbo E, Notredame C. Nextflow enables reproducible computational workflows. Nat Biotechnol. 2017; 35:316–319.

Fridrich A, Modepalli V, Lewandowska M, Aharoni R, Moran Y.) Unravelling the developmental and functional significance of an ancient Argonaute duplication. Nat Commun. 2020;11(1):6187.

Gerber AP, Luschnig S, Krasnow MA, Brown PO, Herschlag D. Genome-wide identification of mRNAs associated with the translational regulator PUMILIO in Drosophila melanogaster. Proc Natl Acad Sci USA. 2006;103(12):4487–92.

Gilbert E, Craggs J, Modepalli V. Gene Regulatory Network that Shaped the Evolution of Larval Apical Organ in Cnidaria. Mol Biol Evol. 2024;41: msad285.

Grasso L, Maindonald J, Rudd S, Hayward D, Saint R, Miller DJ, Ball EE. Microarray analysis provides candidate genes for key roles in coral development. BMC Genomics. 2008;9:540.

Grasso LC, Negri AP, Foret S, Hayward DC, Miller DJ, Ball EE. The biology of coral metamorphosis: molecular responses of coral larvae to inducers of settlement and metamorphosis. Dev Biol. 2011;353:411–419.

Gu Z, Hübschmann D. Make Interactive Complex Heatmaps in R. Bioinformatics. 2022;38:1460–1462. 10.1093/bioinformatics/btab806.

Hadfield MG. Biofilms and Marine Invertebrate Larvae: What Bacteria Produce That Larvae Use to Choose Settlement Sites. Ann Rev Marine Sci. 2011;3:453–470.

Hadfield MG. Why and how marine-invertebrate larvae metamorphose so fast. Sem Cell Dev Biol. 2000;11(6):437–443.

Hayward DC, Miller DJ, Ball EE. Snail expression during embryonic development of the coral *Acropora*: Blurring the diploblast / triploblast divide? Development Genes and Evolution. 2004;214:257–260.

Hayward DC, Samuel G, Pontynen PC, Catmull J, Saint RB, Miller DJ, Ball EE. Localized expression of a DPP/BMP2-4 ortholog in a coral embryo. Proc Natl Acad Sci USA. 2002;99:8106–8111.

Hayward DC, Hetherington S, Behm CA, Grasso L, Foret S, Miller DJ, Ball EE. Differential gene expression at coral settlement and metamorphosis – a subtractive hybridization study. PLoS One. 2011;6: e26411.

Hayward DC, Grasso LC, Saint RB, Miller DJ, Ball EE. The organizer in evolution: Gastrulation and organizer gene expression highlight the importance of Brachyury during development of the coral *Acropora millepora*. Dev Biol. 2015;399:227–247.

He S, Rangel-Huerta E, Hill E, Ellington L, Chen S, Robb S, Majerová E, Drury C, Gibson MC. An evolutionarily conserved Hox-Gbx segmentation code in the rice coral, *Montipora capitata*. bioRxiv preprint. 2024; doi: 10.1101/2024.09.29.615694.

Herchenröther A, Wunderlich TM, Lan J, Hake SB. Spotlight on histone H2A variants: From B to X to Z. Sem Cell Dev Biol. 2023;135:3–12.

Huffmyer AS, Wong KH, Becker DM, Strand E, Mass T, Putnam HM. Shifts and critical periods in coral metabolism reveal energetic vulnerability during development. Curr Biol. 2025;35:1–14;doi 10.1016/j.cub.2025.05.013.

Ishii Y, Hatta M, Deguchi R, Kawata M, Maruyama S. Gene expression alterations from reversible to irreversible stages during coral metamorphosis. Zool Lett. 2022; 8(1):4.

Iwao K, Fujisawa T, Hatta M. A cnidarian neuropeptide of the GLWamide family induces metamorphosis of reef-building corals in the genus Acropora. Coral Reefs. 2002;21:127–129.

Jessop P, Ruzov A, Gering M. Developmental Functions of the Dynamic DNA Methylome and Hydroxymethylome in the Mouse and Zebrafish: Similarities and Differences. Front Cell Dev Biol. 2018;20(6):27.

Jones P, Binns D, Chang HY, Fraser M, Li W, McAnulla C, McWilliam H, Maslen J, Mitchell A, Nuka G, Pesseat S. InterProScan 5: genome-scale protein function classification. Bioinformatics. 2014;30(9):1236–40.

Kitamura M, Koyama T, Nakano Y, Uemura D. Characterization of a natural inducer of coral larval metamorphosis. J Exp Mar Biol Ecol. 2007;340:96–102.

Kitamura M, Schupp PJ, Nakano Y, Uemura D. Luminaolide, a novel metamorphosis-enhancing macrodiolide for scleractinian coral larvae from crustose coralline algae. Tetrahedron Letts. 2009;50:6606–6609.

Langfelder P, Horvath S. WGCNA: an R package for weighted correlation network analysis. BMC Bioinformatics. 2008;9:559. 10.1186/1471-2105-9-559.

Langmead B, Salzberg SL. Fast gapped-read alignment with Bowtie 2. Nat Methods. 2012;9:357–359.

Levy S, Elek A, Grau-Bové X, Menéndez-Bravo S, Iglesias M, Tanay A, Mass T, Sebé-Pedrós A. A stony coral cell atlas illuminates the molecular and cellular basis of coral symbiosis, calcification, and immunity. Cell. 2021;184(11):2973–2987.

Li B, Dewey CN. RSEM: accurate transcript quantification from RNA-Seq data with or without a reference genome. BMC Bioinformatics. 2011;12:323.

Lorković ZJ, Berger F. Heterochromatin and DNA damage repair: Use different histone variants and relax. Nucleus. 2017;8(6):583–588.

Love MI, Huber W, Anders S. Moderated estimation of fold change and dispersion for RNA-seq data with DESeq2. Genome Biol. 2014;15:550.

Mansour TA, Rosenthal JJ, Brown CT, Roberson LM. Transcriptome of the Caribbean stony coral Porites astreoides from three developmental stages. Gigascience. 2016;5(1):33.

Mason BM, Cohen JH. Long-wavelength photosensitivity in coral planula larvae. Biol Bull. 2012;222(2):88–92.

Mason B, Cooke I, Moya A, Augustin R, Lin M-F, Satoh N, Bosch TCG, Bourne DG, Hayward DC, Andrade N, Forêt S, Ying H, Ball EE, Miller DJ. AmAMP1 from *Acropora millepora* and damicornin define a family of coral-specific antimicrobial peptides related to the Shk toxins of sea anemones. Dev Comp Immunol. 2021;114: 103866.

Mason B, Hayward DC, Moya A, Cooke I, Sorenson A, Brunner R, Andrade N, Huerlimann R, Bourne DG, Schaeffer P, Grinblat M, Ravasi T, Ueda N, Tang SL, Ball EE, Miller DJ. Microbiome manipulation by corals and other Cnidaria via quorum quenching. Curr Biol. 2024;34(14):3226–3232.e5.

Mason BM, Koyanagi M, Sugihara T, Iwasaki M, Slepak V, Miller DJ, Sakai Y, Terakita A. Multiple opsins in a reef-building coral, *Acropora millepora*. Scientific Reports 2023;23:1628.

Matsumoto N, Sato K, Nishimasu H, Namba Y, Miyakubi K, Dohmae N, Ishitani R, Siomi H, Siomi MC, Nureki O. Crystal structure and activity of the endoribonuclease domain of the piRNA pathway factor Maelstrom. Cell Reps. 2015;11:366–375.

Mazza ME, Pang K, Reitzel AM, Martindale MQ, Finnerty JR. A conserved cluster of three PRD-class homeobox genes (homeobrain, rx and orthopedia) in the Cnidaria and Protostomia. EvoDevo. 2010;1:1–15.

Meyer E, Aglyamova GV, Matz MV. Profiling gene expression responses of coral larvae (Acropora millepora) to elevated temperature and settlement inducers using a novel RNA-Seq procedure. Mol Ecol. 2011;20:3599–616.

Meyer E, Aglyamova GV, Wang S, Buchanan-Carter J, Abrego D, Colbourne JK, Willis BL, Matz MV. Sequencing and de novo analysis of a coral larval transcriptome using 454 GSFlx. BMC Genomics. 2009;10:1–8.

Michail C, Rodrigues Lima F, Viguier M, Deshayes F. Structure and function of the lysine methyltransferase SETD2 in cancer: From histones to cytoskeleton. Neoplasia. 2025;59:101090.

Miller DJ, Hemmrich G, Ball EE, Hayward DC, Khalturin K, Funayama N, Agata K, Bosch TCG.The innate immune repertoire in Cnidaria – ancestral complexity and stochastic gene loss. Genome Biol. 2007;8:R59.

Mishima Y, Tomari Y. Pervasive yet nonuniform contributions of Dcp2 and Cnot7 to maternal mRNA clearance in zebrafish. Genes Cells. 2017;22(7):670–678.

Modepalli V, Fridrich A, Agron M, Moran Y. The methyltransferase HEN1 is required in Nematostella vectensis for microRNA and piRNA stability as well as larval metamorphosis. PLoS Genet. 2018;14(8):e1007590.

Mohr F, Döhner K, Buske C, Rawat VP. TET genes: new players in DNA demethylation and important determinants for stemness. Exp Hematol. 2011; 39(3):272–81.

Moran Y, Agron M, Praher D, Technau U. The evolutionary origin of plant and animal microRNAs. Nat Ecol Evol. 2017;1(3):27.

Morse DE, Hooker N, Morse AN, Jensen RA. Control of larval metamorphosis and recruitment in sympatric agariciid corals. J Exp Mar Biol Ecol. 1988;116(3):193–217.

Negri AP, Webster NS, Hill RT, Heyward AJ. Metamorphosis of broadcast spawning corals in response to bacteria isolated from crustose algae. Mar Ecol Prog Ser. 2001;223:121–131.

Okubo N, Hayward DC, Forêt S, Ball EE. A comparative view of early development in the corals *Favia lizardensis*, *Ctenactis echinata*, and *Acropora millepora* - morphology, transcriptome, and developmental gene expression. BMC Evol Biol. 2016;16:1–12.

Okubo N, Mezaki T, Nozawa Y, Nakano Y, Lien YT, Fukami H, Hayward DC, Ball EE Comparative embryology of eleven species of stony corals (Scleractinia). PLoS One. 2013;8(12): e84115.

Osakabe A, Molaro A Histone renegades: Unusual H2A histone variants in plants and animals. Sem Cell Dev Biol. 2023;135:35–42.

Palmer CV, Modi CK, Mydlarz LD. Coral fluorescent proteins as antioxidants. PLoS One. 2009;4(10):e7298.

Praher D, Zimmermann B, Genikhovich G, Columbus-Shenkar Y, Modepalli V, Aharoni R, Moran Y, Technau U. Characterization of the piRNA pathway during development of the sea anemone Nematostella vectensis. RNA Biol. 2017; 14(12):1727–1741.

Quinlan ZA, Bennett MJ, Arts MGI, Levenstein M, Flores D, Tholen HM, Tichy L, Juarez G, Haas AF, Chamberland VF, Latijnhouwers KRW, Vermeij MJA, Johnson AW, Marhaver KL, Kelly LW. Coral larval settlement induction using tissue-associated and exuded coralline algae metabolites and the identification of putative chemical cues. Proc Roy Soc B. 2023;290:20231476. 10.1098/rspb.2023.1476.

Rabani M, Pieper L, Chew G-L, Schier AH. A Massively Parallel Reporter Assay of 3’ UTR Sequences Identifies In Vivo Rules for mRNA Degradation. Mol Cell. 2017; 68(6):1083–1094.e5.

Reyes-Bermudez A, Desalvo MK, Voolstra CR, Sunagawa S, Szmant AM, Iglesias-Prieto R, Medina M. Gene expression microarray analysis encompassing metamorphosis and the onset of calcification in the scleractinian coral Montastraea faveolata. Mar Genomics 2009;2:149–159.

Ramon-Mateu J, Ferraioli A, Teixidó N, Domart-Coulon I, Houliston E, Copley RR. Aboral cell types of Clytia and coral larvae have shared features and link taurine to the regulation of settlement. Science Adv. 2025;16;11(20):eadv1159.

Reyes-Bermudez A, Villar-Briones A, Ramirez-Portilla C, Hidaka M, Mikheyev AS Developmental Progression in the Coral *Acropora digitifera* Is Controlled by Differential Expression of Distinct Regulatory Gene Networks. Genome Biol Evol. 2016;8(3):851–70.

Salih A, Larkum A, Cox G, Kuhl M, Hoegh-Guldberg Fluorescent pigments in corals are photoprotective. Nature.2000;408:850–853.

Shoguchi E, Beedessee G, Hisata K, Tada I, Narisoko H, Satoh N, Kawachi M, Shinzato C. A New Dinoflagellate Genome Illuminates a Conserved Gene Cluster Involved in Sunscreen Biosynthesis. Genome Biol Evol. 2021;13:evaa235.

Sato K, Siomi MC. Functional and structural insights into the piRNA factor Maelstrom. FEBS Lett. 2015;589:1688–1693.

Siboni N, Abrego D, Puill-Stephan E, King WL, Bourne DG, Raina J-B, Seymour JR, Harder T. Crustose coralline algae that promote coral larval settlement harbor distinct surface bacterial communities. Coral Reefs 2020;39:1703–1713.

Spike CA, Tsukamoto T, Greenstein D. Ubiquitin ligases and a processive proteasome facilitate protein clearance during the oocyte-to-embryo transition in *Caenorhabditis elegans*. Genetics. 2022;221(1):iyac051.

Strader ME, Davies SW, Matz MV (2015) Differential responses of coral larvae to the colour of ambient light guide them to suitable settlement microhabitat. R Soc Open Sci 2(10):150358

Strader ME, Aglyamova GV, Matz MV. Molecular characterization of larval development from fertilization to metamorphosis in a reef-building coral. BMC Genomics. 2018;19(1):1–17.

Takeuchi T, Yamada L, Shinzato C, Sawada H, Satoh N. Stepwise Evolution of Coral Biomineralization Revealed with Genome-Wide Proteomics and Transcriptomics. PLoS One 2016;11(6):e0156424.

Tebben J, Motti CA, Siboni N, Tapiolas DM, Negri AP, Schupp PJ, Kitamura M, Hatta M, Steinberg PD, Harder T. Chemical mediation of coral larval settlement by crustose coralline algae. Sci Rep. 2015;5:10803.

Technau U. Gastrulation and germ layer formation in the sea anemone Nematostella vectensis and other cnidarians. Mech Dev. 2020;163:103628.

Török A, Schiffer PH, Schnitzler CE, Ford K, Mullikin JC, Baxevanis AD, Bacic A, Frank U, Gornik SG. The cnidarian Hydractinia echinata employs canonical and highly adapted histones to pack its DNA. Epigenetics Chromatin. 2016;9(1):1–17.

Tran C, Hadfield MG. Are G-protein-coupled receptors involved in mediating larval settlement and metamorphosis of coral planulae? Biol Bull 2012;222(2):128–36.

Turnlund AC, Vanwonterghem I, Botté ES, Randall CJ, Giuliano C, Kam L, Bell S, O’Brien P, Negri AP, Webster NS, Lurgi M. Linking differences in microbial network structure with changes in coral larval settlement. ISME Comm. 2023;3:114.

Vastenhouw NL, Cao WX, Lipshitz HD. The maternal-to-zygotic transition revisited. Development. 2019;146(11): dev161471

Walker NS, Fernández R, Sneed JM, Paul VJ, Giribet G, Combosch DJ. Differential gene expression during substrate probing in larvae of the Caribbean coral *Porites astreoides*. Mol Ecol. 2019;28(22):4899–4913.

Wickham H. Data Analysis, in: Wickham, H. (Ed.), Ggplot2: Elegant Graphics for Data Analysis. Springer International Publishing, Cham, 2016; pp. 189–201. 10.1007/978-3-319-24277-4_9.

Wu X, Zhang Y. TET-mediated active DNA demethylation: mechanism, function and beyond. Nat Rev Genet. 2017;18(9):517–534.

Yasuoka Y, Shinzato C, Satoh N. The Mesoderm-Forming Gene brachyury Regulates Ectoderm-Endoderm Demarcation in the Coral Acropora digitifera. Curr Biol. 2016;26(21):2885–2892.

