## Supplementary figures and images for "Major transitions in early coral development: novel insights enabled by visualisation of a comprehensive transcriptomic dataset for *Acropora millepora*"

### Supplentary data

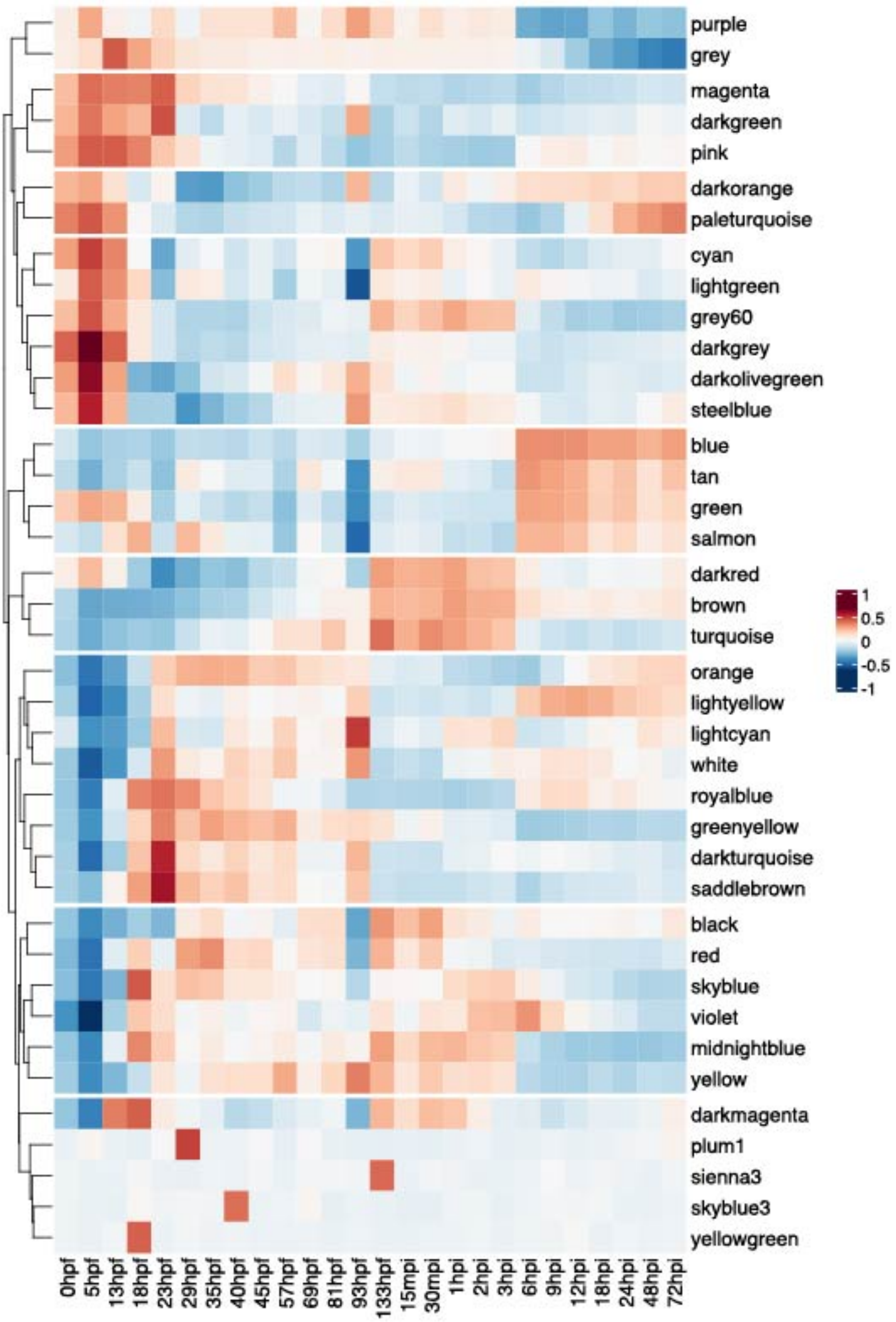

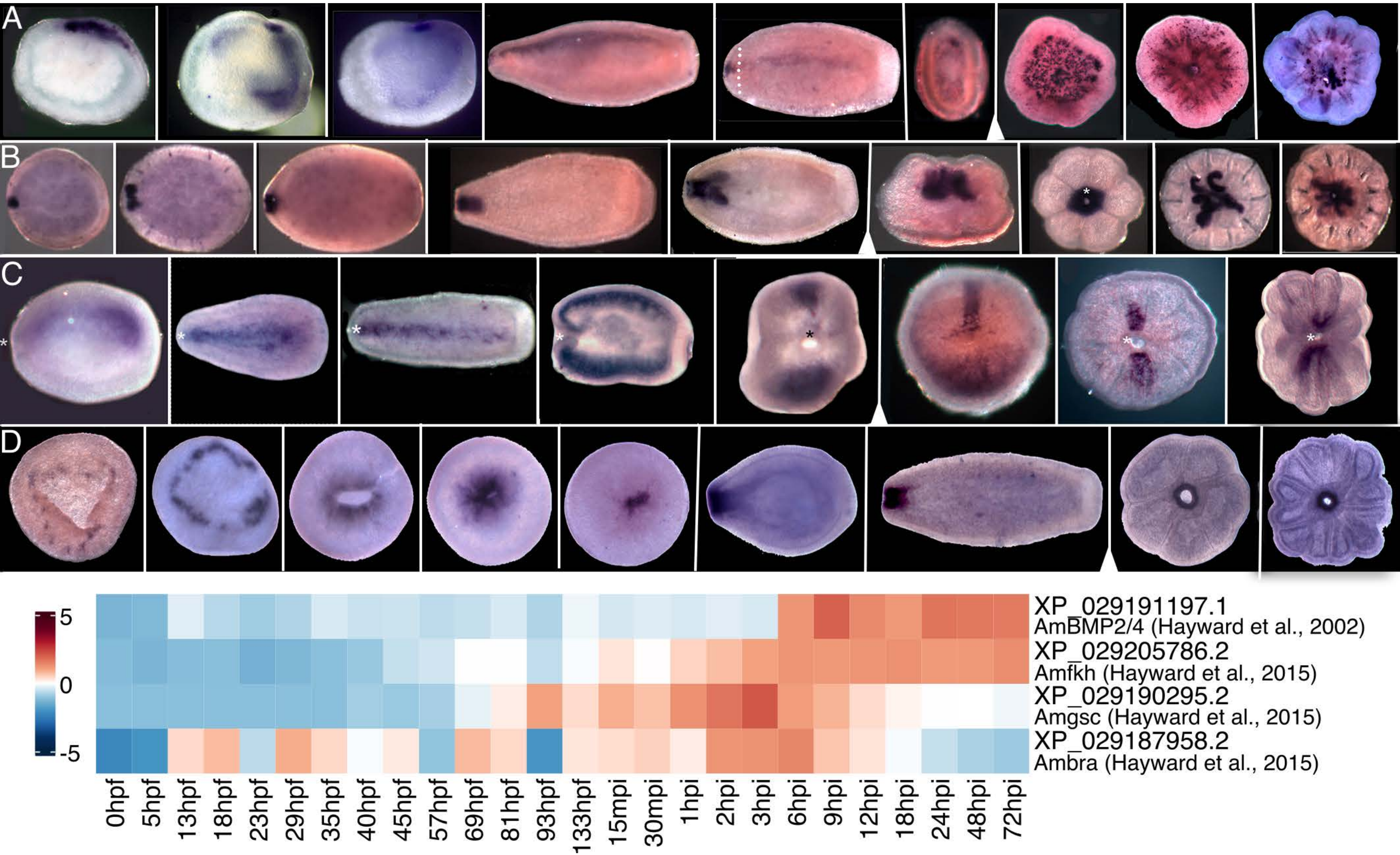

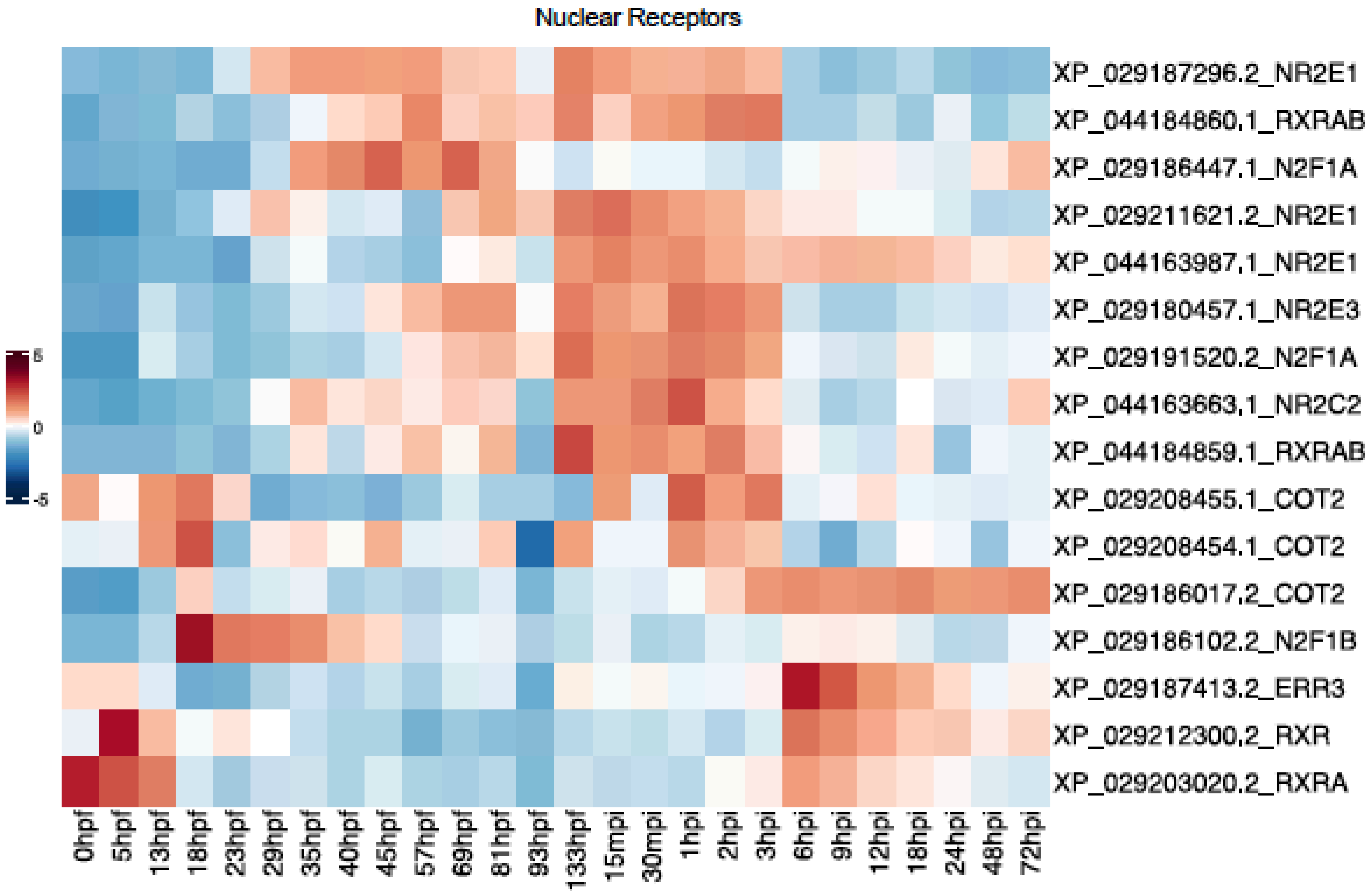
